# Ultrafast Light Targeting for High-Throughput Precise Control of Neuronal Networks

**DOI:** 10.1101/2021.06.14.448315

**Authors:** Giulia Faini, Clément Molinier, Cécile Telliez, Christophe Tourain, Benoît C. Forget, Emiliano Ronzitti, Valentina Emiliani

**Affiliations:** Institut de la Vision, Sorbonne Université, Inserm S968, CNRS UMR7210, 17 Rue Moreau, 75012 Paris

## Abstract

Understanding how specific sets of neurons fire and wire together during cognitive-relevant activity is one of the most pressing questions in neuroscience. Two-photon, single-cell resolution optogenetics based on holographic light-targeting approaches enables accurate spatio-temporal control of individual or multiple neurons. Yet, currently, the ability to drive asynchronous activity in distinct cells is critically limited to a few milliseconds and the achievable number of targets to several dozens. In order to expand the capability of single-cell optogenetics, we introduce an approach capable of ultra-fast sequential light targeting (FLiT), based on switching temporally focused beams between holograms at kHz rates. We demonstrate serial-parallel photostimulation strategies capable of multi-cell sub-millisecond temporal control and many-fold expansion of the number of activated cells. This approach will be important for experiments that require rapid and precise cell stimulation with defined spatio-temporal activity patterns and optical control of large neuronal ensembles.

## INTRODUCTION

Optogenetic neuronal excitation using single-photon widefield illumination has already proven its enormous potential in neuroscience, enabling the optical manipulation of entire neuronal networks and to disentangle their role in the control of specific behaviors^1,2^. However, establishing how the activity of a single neuron or neuronal ensemble impacts a specific behavior, or how functionally identical neurons are connected and involved in a particular task, requires the precise control of single or multiple cells independently in space and time. This has imposed a transition from widefield optogenetics into a more sophisticated technology that we termed few years ago: circuit optogenetics^3^. Circuit optogenetics combines progress in opsin engineering, holographic light shaping and high-power fiber laser development. Using two-photon holographic illumination of fast-photocycle-, soma-targeted-opsins, it permits single or multi-spike generation with cellular resolution, sub-millisecond precision and high spiking rates deep in tissue^3^. Using multiplexed spiral scanning^4^ or multiplexed temporally focused light shaping approaches^5–7^, combined with high energy fiber lasers and soma-targeted opsins^8,9^, it also enables simultaneous control of multiple targets in 3D ∼*mm*^3^ volumes at cellular resolution^10–12^.

The unprecedented spatiotemporal precision of circuit optogenetics has enabled high throughput connectivity mapping in living zebrafish larvae^13^ and probing rod bipolar cell output across multiple layers of the mouse retina^14^. Combined with two photon Ca^2+^ imaging and behavioural assays, circuit optogenetics has been used to show that the activation of few cells can bias behavior by triggering the activity of precisely-defined ensembles in the mouse visual cortex^11,12,15^. Importantly, sequential projection of multiple holographic patterns at variable time intervals in the mouse olfactory bulb has revealed how the perceptual responses of mice not only depend on the specific group of cells and cell numbers activated but also on their relative activation latency^16^.

These pioneering works suggest a number of new exciting experimental paradigms for circuit optogenetics, e.g., the investigation of the temporal bounds of functional connectivity within which neurons “fire and wire together”, or how many targets need to be activated to perturb complex behavioral responses or how large neuronal ensembles, eventually spanning across multiple cortical layers, are functionally connected. Answering these questions requires the capability to manipulate neuronal activity at fine (sub-millisecond) temporal scales and/or large cell populations, which ultimately requires overcoming the current intrinsic technological limitations of holographic light patterning, specifically the low speed of liquid crystal spatial light modulators (LC-SLMs) and the high illumination power necessary for multi-target excitation.

Multi-target optogenetics uses holographic light shaping to multiplex the excitation beam to multiple locations, combined with either spiral scanning or temporally focused light shaping approaches^17,18^.

In spiral scanning approaches, a LC-SLM is used to multiplex the illumination beam into several diffraction limited spots which are scanned in spiral trajectories using a pair of galvanometric mirrors (GM), each spanning a different neuron.

Several multiplexed temporally focused light shaping (MTF-LS) approaches have been developed^5^, which differ in terms of the approach used for light patterning. Generally, MTF-LS systems are comprised of three units: (1) a beam shaping unit sculpts light into particular forms, (2) a diffraction grating placed in a conjugate image plane confines photostimulation to a shallow axial region with cellular dimensions and (3) a LC-SLM multiplexes the sculpted light to multiple sample locations (Fig.1A).

This configuration admits multiple variants depending on the beam shaping unit used, which also defines the extent of the beam profile at the multiplexing LC-SLM^5^. Beam shaping units based on computer generated holography illuminate the second LC-SLM with either a single chirped hologram of the size of the LC-SLM matrix^6^ or with multiple chirped holograms^19^, while the use of an expanded gaussian beam^20^ or of the generalized phase contrast method^6^, produces a chirped horizontal line (typically ∼ 2 mm high and ∼ 16 mm wide) covering the horizontal dimension of the LC-SLM or a diffused chirped spot which illuminates the entire LC-SLM matrix^7^ (Supplementary Fig.1). In all described approaches, sequential generation of independent illumination patterns is achieved by projecting multiple holograms at a rate limited by the LC-SLM refresh rate (60-500 Hz) and cell illumination times (from ∼1 ms to dozens of ms).

Moreover, multi-target illumination based on holography requires powerful lasers since the laser energy is divided between different targets^3,17,21^. While in principle this enables the simultaneous photostimulation of hundreds of target cells, the necessity of maintaining brain temperature within physiological thresholds^10,22,23^ has thus far limited the maximum number to a few dozens.

Here, we suggest a new approach for ultra-fast sequential light targeting (FLiT) based on rapid displacement of temporally-focused sculpted light through multiple, vertically-aligned, holograms. We demonstrated that optogenetic FLiT enables tuning of distinct cells with microsecond resolution and a 20 (or more) times increase of achievable targets with minimal thermal crosstalk.

This novel capability of arbitrarily desynchronizing (or synchronizing) groups of neurons will facilitate the study of the influence of spike timing on synaptic integration, plasticity and information coding, and scaling up the number of cells activable whilst remaining safely below the threshold for thermal damage.

## RESULTS

### Ultra Fast Light-Targeting (FliT)

Here we introduce a new configuration of MTF-LS for ultra-fast sequential light targeting (FLiT), where the multiplexing LC-SLM is addressed with multiple vertically tiled holograms. A galvanometric mirror (GM) is incorporated upstream (Fig.1B) to sweep the chirped expanded gaussian beam across the holograms and generate sequential 2D (Fig.1C; Supplementary Movie 1) or 3D (Fig.1D) illumination patterns.

**Figure 1:**
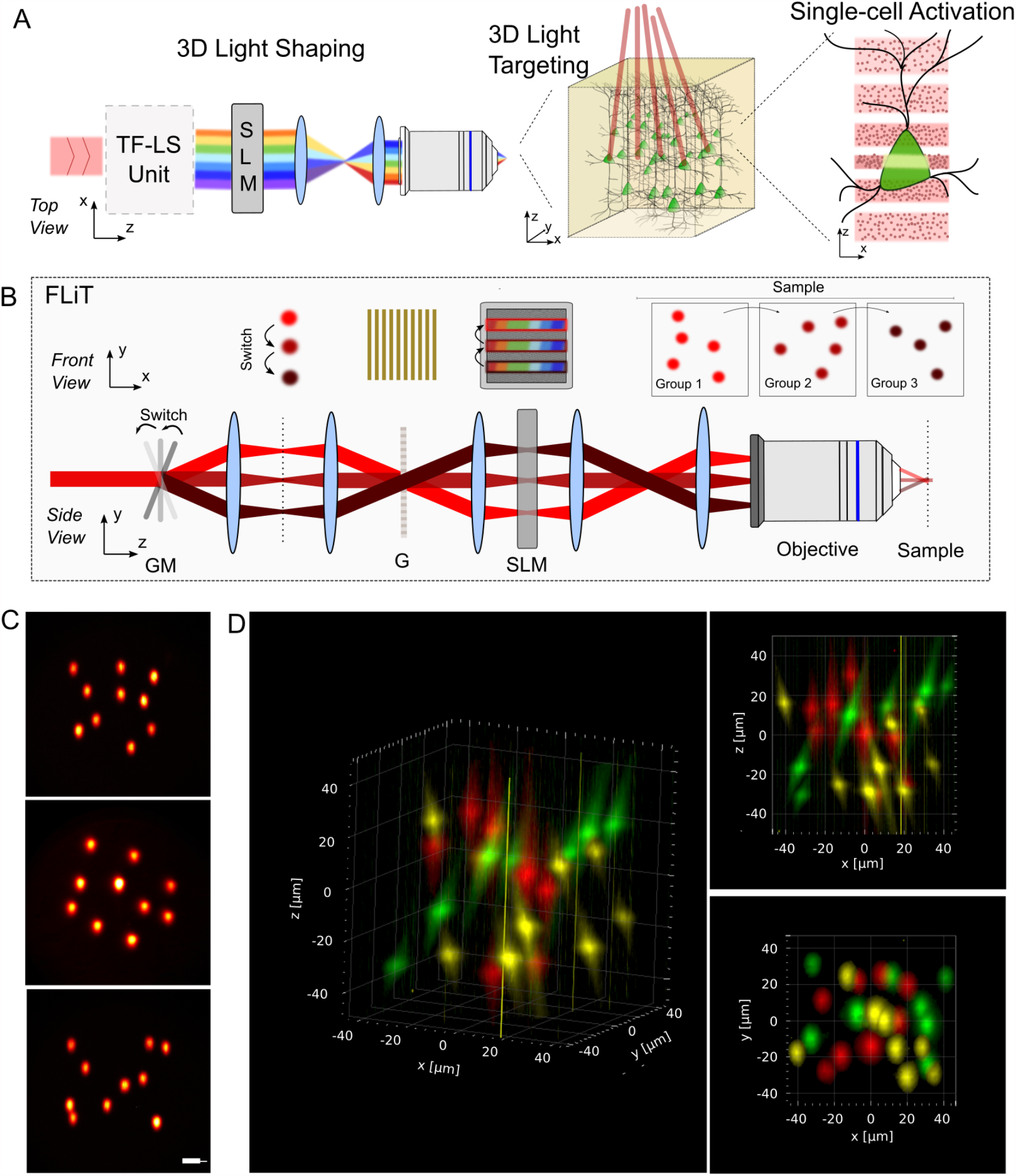
FLiT optical characterization. **(A)** General optical scheme for temporally focused light shaping. A first light-shaping temporally focusing architecture (LS-TF) allows *(i)* sculpting light into particular forms and *(ii)* temporally focusing the photons to confine photostimulation to a shallow axial region with cellular dimensions. A subsequent LC-SLM modulation allows multiplexing the sculpted light to multiple 3D sample locations. **(B)** Optical setup of FLiT. A pulsed collimated beam (red line) is reflected by a galvanometric mirror (GM) onto a diffracting grating (G) via a *4f*-telescope. Diffracted off the grating, the beam is collimated onto a liquid crystal spatial light modulator (LC-SLM) in the form of a horizontal (i.e., orthogonal to the orientation of the grating lines) spatially chirped strip of light. The LC-SLM is imaged onto the back aperture of an objective lens so that *ad hoc* phase-modulation on the LC-SLM allows multiplexing the initial beam and generating a multi-site temporally focused pattern of light in the sample. As deflection of the beam by the GM results into a translation of the illuminating bands on the LC-SLM (dark red lines), addressing the LC-SLM with *n* independent tiled holograms *φ*_*i*_ leads to fast switch of different groups of light patterns into the sample. The top and bottom drawing represents the XY and the YZ plane views, respectively. **(C)** x-y 2PE fluorescence cross-sections of different groups of randomly distributed spots generated in the sample focal plane by addressing the *i-th* hologram *φ*_*i*_ of an LC-SLM subdivided in 20 tiled holograms: hologram *φ*_*8*_ (Top), hologram *φ*_*10*_ (Middle) and hologram *φ*_*12*_ (Bottom). Scale bar: 20µm. **(D)** 2PE fluorescence of different groups of spots generated by different tiled holograms *φ*_*i*_ randomly distributed across a 120×120×70 µm^3^ volume. Different colors correspond to different tiled hologram (hologram *φ*_*4*_, yellow; hologram *φ*_*10*_, red; and hologram *φ*_*16*_, green).

To characterize the optical properties of FliT, we first characterized the effect of the hologram tiling on the holographic spot intensity, ellipticity (i.e., ratio between vertical (y) and horizontal (x) length) and axial resolution by projecting on a thin rhodamine layer a single spot encoded by tiled holograms with different vertical extents (see Methods). We did not observe a significant deterioration of the axial resolution decreasing the tile extent to 12 lines (corresponding to 50 tiled holograms for the particular LC-SLM used), while we observed a decrease of a factor ≥2 in spot intensity and ellipticity by decreasing the tile extent to ≤20 lines (corresponding to 30 tiled holograms) (Supplementary Fig.2).

We then evaluated, for the case of 30 lines per tiled hologram (20 tiled phase holograms, *φ*_*i*_), the influence of the location of the hologram on the LC-SLM on spot intensity and axial resolution by deflecting the chirped beam across 20 2D holograms (each encoding the same group of spots) using the GM. For each hologram, the intensity was homogeneously distributed among spots generated in a field of excitation (FoE) of 120 × 120 μm^2^ (Fig.1C; Supplementary Fig.3). For patterns encoded in holograms located in central regions of the LC-SLM, we observed a ≥25% higher average intensity than in patterns generated by distal holograms (*φ*_*i*_ ≤4; *φ*_*i*_ ≥14; Supplementary Fig.3). Although we observed a moderate axial tilt of the spots generated by using distal holograms, the axial resolution of the spots was preserved both within the FoE and while scanning across the different holograms (6.5 ± 0.5 µm; Supplementary Fig.4). Importantly, spot intensity homogeneity and axial confinement were maintained when spots were randomly distributed in 120 × 120 × 70 µm^3^ volume (Supplementary Fig.5).

Next, we studied how the velocity of the scan unit defines the temporal resolution of FLiT. Specifically, we tested the minimum switching time to *(i)* move between two adjacent holograms (Fig.2A-B) and *(ii)* sequentially illuminate all holograms at constant rate (Fig.2E-F). For this, we generated 20 equivalent holograms each projecting a single spot on a photodiode placed in a conjugate image plane (Fig2A and Fig.2E) (see details in Methods). In the first case, we measured a switching time of 90 ± 10 μs (Fig.2C-D, n = 30 measurements), while in the second case, we could reach a switching time of 50 ± 10 μs (Fig.2G-H, n = 30 measurements).

**Figure 2:**
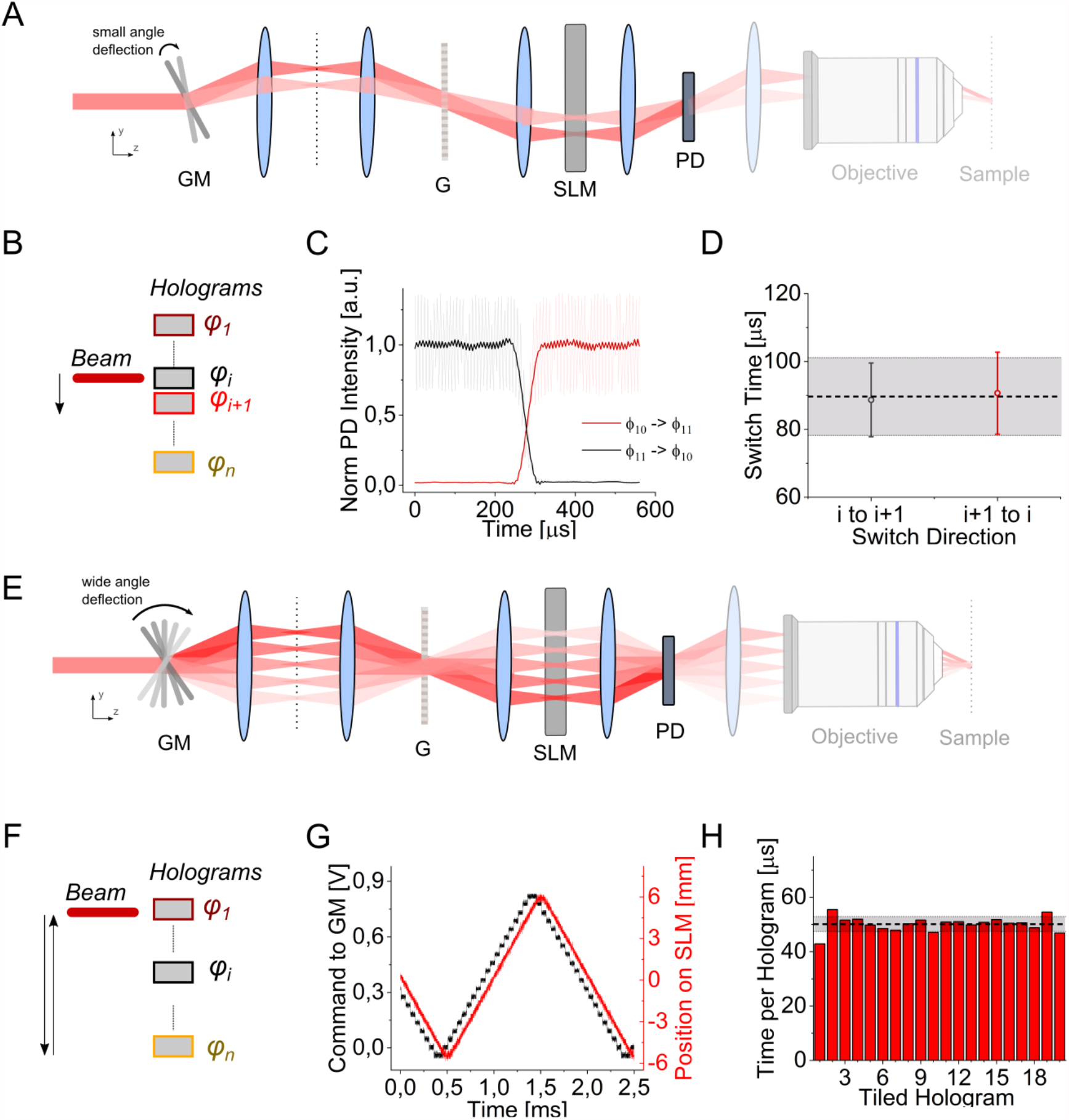
FLiT switching time. (**A**) Switching time between two adjacent tiled holograms (*φ*_*i*_ and *φ*_*i*+*1*_) is measured by means of a photodiode (PD) placed in an image conjugate plane while driving the galvanometric mirror (GM) with small angles single-step voltage inputs. (**B**) Scheme of illumination switch between tiled hologram *φ*_*i*_ and *φ*_*i*+*1*_ on the SLM display corresponding to sequence depicted in (A). (**C**) Representative intensity response of the PD when GM is switched from hologram *φ*_*11*_ (encoding for an individual spot in the middle of PD) to hologram *φ*_*10*_ (deviating the beam out of the PD) (*black line*) or, vice versa, from hologram *φ*_*10*_ (encoding for an individual spot in the middle of PD) to hologram *φ*_*11*_ (deviating the beam out of the PD) (*red line*). (**D**) Switch time calculated as the time taken for the signal to rise/fall between 3% and 97% of the maximum intensity, when the spot is encoded in hologram *φ*_*i*_ and GM is switched from hologram *φ*_*i*_ to *φ*_*i*+*1*_ (black symbols) or vice versa (red symbols). Horizontal black line and grey bands indicate the global mean and SD switching time, respectively. (**E**) Switch time to sequentially illuminate all holograms at constant rate from *φ*_*1*_ to *φ*_*n*_ is measured by driving the galvanometric mirror (GM) with a wide-angle staircase voltage input. (**F**) Scheme of the illumination switch to sequentially illuminate all holograms from *φ*_*1*_ to *φ*_*n*_ on the SLM display corresponding to the sequence depicted in (E). (**G**) GM voltage input (black line) and corresponding position of the incoming beam on the LC-SLM (red line) when GM is driven as depicted in (E). (**H**) Dwell time of each hologram *φ*_*i*_ of the LC-SLM while GM is driven as depicted in (E) and *φ*_*i*_ only encodes an individual spot in the middle of the PD. Horizontal black and grey lines indicate the mean and SD dwell-time over all holograms. For all experiments the LC-SLM was dived in 20 tiled holograms.

Taken together these results indicate that FLiT enables sequential generation of multiple patterns with no significant deterioration of spot quality or axial resolution for up to 12 phase holograms. By scanning the chirped beam among the multiple holograms, it is possible to reach up to few tens of kHz switching rate, which is more than one order of magnitude faster than what is achievable with alternative parallel approaches using phase modulation^11^.

### Precisely replaying physiological patterns of activity

In order to demonstrate the capabilities of FLiT to control neuronal activity at high switching rate, we photoactivated neurons expressing the soma-restricted opsin ST-ChroME^10^ while recording cellular activity via whole-cell patch clamp recordings in acute cortical brain slices (see Methods and Supplementary Fig.6A).

We initially tested the illumination conditions (excitation power, illumination time and spot size) for reliable action potential (AP) generation. Consistent with results previously obtained in standard 2P holographic configurations^9,21,24^, APs could be reliably elicited with sub-ms jitter (0.25 ± 0.13 ms; n= 13 cells; Supplementary Fig.7) upon selective targeting of cell somata (spot size = 10 µm) with illumination times as short as 4-5 ms. These values and the LC-SLM refresh rate (60-500 Hz)^10,11^ set the effective temporal resolution for sequential photostimulation in common holographic light patterning techniques.

Here, we demonstrate that FLiT overcomes this limit by enabling sub-millisecond temporal resolution independently of the illumination time and LC-SLM switching rate (Fig.3A). Briefly, if two groups of cells (group A and B) need to be activated with a temporal delay shorter than the necessary illumination time (dwell-time), one can use 3 phase holograms: the first one (φ_A_) generating a light pattern to excite the group A, the second one (φ_B_) to excite the group B and the intermediate one (φA_B_) to excite both groups, A and B. By steering the beam across the three holograms, each with a specific illumination time and intensity, it is possible to sequentially stimulate the two groups of cells with tightly controlled delays, only limited by the GM scanning time (i.e., in our case ≥90 μs). Notably, the same principle can be extended to *n* groups of cells by using *2n-1* tiled holograms and sequentially addressing the different groups in parallel or individually (Supplementary Fig.8). We call this configuration serial/parallel FLiT (S/P-FLiT).

We demonstrated the capability of S/P-FLiT for ultra-fast sequential light targeting by photostimulating two ST-ChroME-expressing neurons while monitoring the evoked activity by double-patch electrophysiological recordings (Fig.3B). First, we verified that amplitudes and kinetics of induced photocurrents were not affected by switching the illumination between the different holograms (Supplementary Fig.9). We then assessed the precision in controlling the relative spiking time among the two cells by photoactivating the two patched neurons with tightly controlled delays, *δt*, ranging from 0.2 to 3 ms, while measuring the corresponding spiking delay time 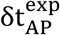 (Fig.3B). We found that spike delays 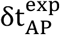 can be controlled with few hundreds µs temporal accuracy, 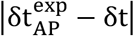, (96 ± 114 μs, n = 12 pairs of cells; Fig.3C and Supplementary Fig.10). Furthermore, the photocurrent magnitude was found to be approximately independent of the vertical position of the tiled hologram (Supplementary Fig.11).

**Figure 3:**
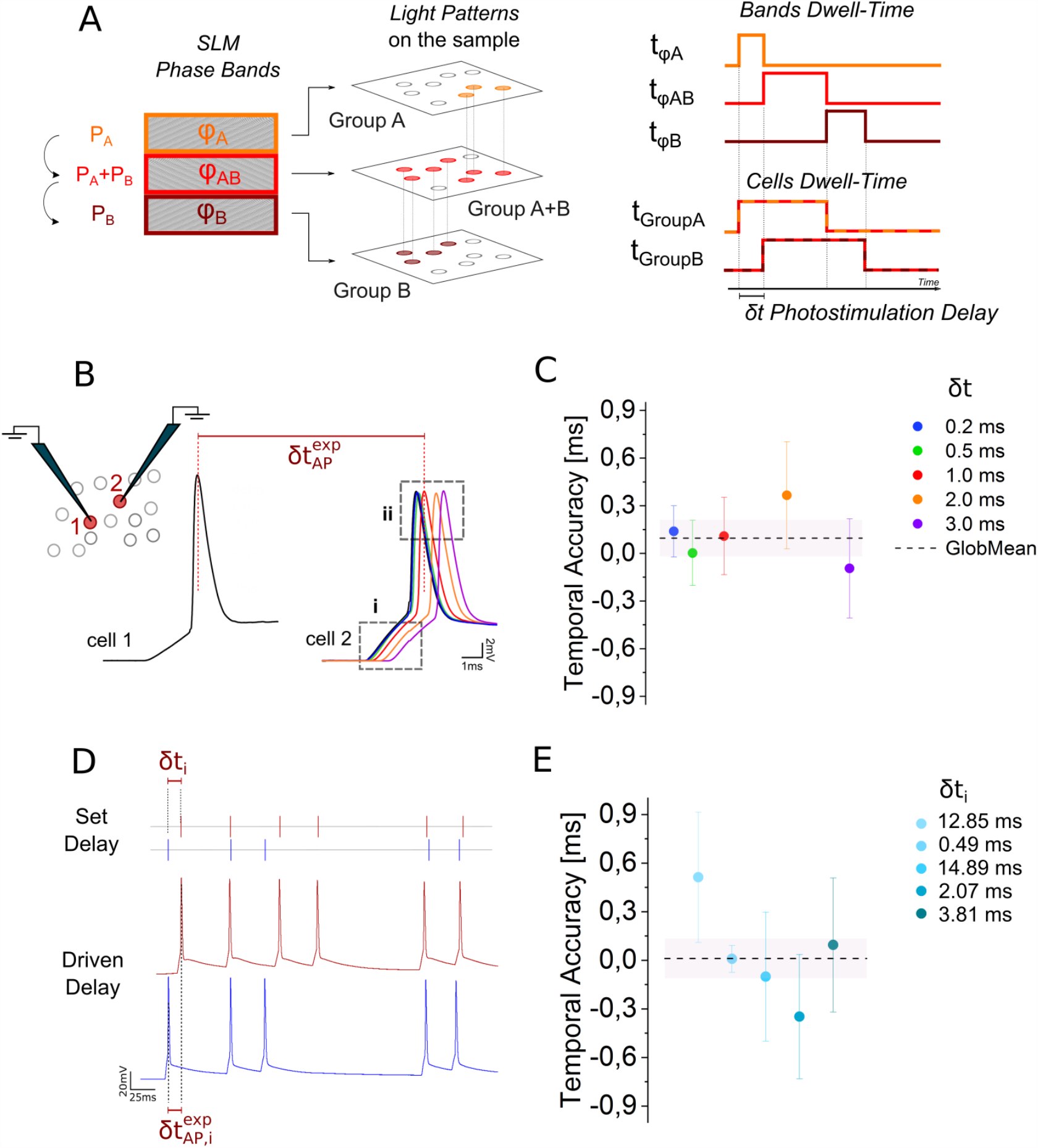
Tuning control of neuronal activity in targeted neurons by S/P-FLiT. **(A)** Conceptual scheme of S/P FliT. The LC-SLM is tiled in different regions each encoding different phase masks. In the present example, phase mask *φ*_*A*_ and *φ*_*B*_ encode for group of spots A and B, while phase mask *φ*_*AB*_ encodes for a comprehensive pattern including group A and group B. By steering the beam vertically across the phase masks with predetermined dwell-times and illumination intensities per each mask, it is possible to set arbitrary delays of activation between groups of spots. In the illustred example, the laser dwell-time is t_*φA*_, t_*φAB*_, t_*φB*_, and the illumination power is *P*_*A*_, *P*_*A*_ + *P*_*B*_, *P*_*B*_ on *φ*_*A*_, *φ*_*AB*_, *φ*_*B*_ respectively. Importantly on comprehensive phase mask *φ*_*AB*,_ the distribution of intensity must be computationally set to maintain an amount of power P_A_ and P_B_ on subgroup A and B, respectively. Overall, this scheme yields an activation time t_*φA*_ +t_*φAB*_ for group A, t_*φAB*_ +t_*φB*_ for group B and a delay of activation between group A and group B *δt* equivalent to t_*φA*_. The scheme displayed is meant to represent *n* groups of spots; their number is here limited to 2 for presentation purposes only. **(B)** Representative light-driven APs from two double-patched ST-ChroME-expressing neurons by imposing different delays δt ranging from 0.2ms to 3ms. **(C)** Temporal accuracy calculated as the difference between imposed δt and experimental 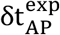 delays, 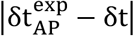. Circle symbols represent different *δt* delays (data are shown as mean ± SD). Horizontal dashed black line and grey bands represents global mean and SD, respectively. Error bars are SD on n = 12 pairs of cells. Mean AP accuracy is 96 ± 114 µs. **(D)** Representative light-driven APs from two double-patched ST-ChroME-expressing neurons (Bottom) by imposing a random spiking patterns (Top) featuring inter-spike time intervals *δt*_*i*_. **(E)** Temporal accuracy calculated as difference between imposed δt_*i*_ and experimental 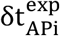 delays of the *i-th* AP pair, 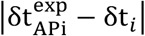. Circles from light to dark blue indicate temporal accuracy from subsequent pairs of APs (data are shown as mean ±SD). Mean temporal accuracy is 11 ± 122 µs (n = 12 pair of cells). Mean photostimulation power is 37.7 ± 21.3 mW. Illumination dwell-time ranges between and 2-5ms. 1030nm illumination has been used.

Finally, we demonstrated the capability of S/P-FLiT to precisely mimic random spiking activity in two distinct neurons. To this end, we photostimulated two neurons with distinct spiking patterns based on physiological activity (Fig.3D). Light-driven mimicking was precisely controlled with few hundreds of µs temporal accuracy, 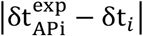, where *i* indicates the AP ordinal number in the train (11 ± 122 µs; n = 12 pairs of cells; Fig.3E and Supplementary Fig.12).

Taken together, these results indicate that S/P-FLiT enables precise sub-millisecond tuning of neuronal activity in distinct neurons or groups of neurons.

### High-throughput activation of multiple cells by tuning Illumination to match properties of opsin photocycle

Here we demonstrated the capability of FliT to scale up the achievable number of targets for parallel multi-cell illumination.

In conventional parallel illumination approaches, the simultaneous excitation of *n* targets requires an excitation power of *n* · *P*_*std*_, where *P*_*std*_ is the excitation power which needs to be continuously applied for a time *t*_*std*_ to activate a single target (Fig.4A). Here we demonstrate that FliT approach enables targeting the same number of cells by using cyclic illumination with μs flashes of light and a factor of 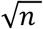 lower power whilst maintaining identical latency and jitter. Alternatively, a factor of 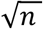 more cells can be targeted using the same amount of power. We called this configuration serial-parallel multi-cell activation FliT (Multi-S/P FliT).

Briefly, steady illumination of a neuron for a time *t*_*std*_ shorter than the opsin rise time generates an exponential increase in photocurrent (Fig.4A) and eventually AP generation. Capitalizing on the properties of the opsin photocycle, a similar photocurrent can be generated by using cyclic illumination consisting of *N*_*cyc*_ short illumination pulses each of duration *t*_*cyc*_ ≪ *t*_*std*_, provided *(i)* the time interval, *T*_*cyc*_, between two pulses is shorter than the off-time decay of the opsin (Supplementary Note 1) and *(ii)* the excitation power, *P*_*cyc*_, generates in a time *t*_*cyc*_ a photocurrent that after a time 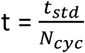, equals the one that a steady illumination with power *P*_*std*_ would generate in the same time. It can be shown that this condition is realized for 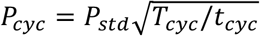 (Supplementary Note 1). If these conditions are satisfied, FLiT can rapidly reposition the excitation light onto 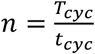 different locations in a time *t* = *T*_*cyc*_, thus enabling quasi-simultaneous activation of *n* targets with a power only 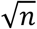 higher than *P*_*std*_ (Fig.4B).

In order to demonstrate this configuration, we divided the LC-SLM in *n* tiled holograms, encoding a soma-targeted illumination of a patched ST-ChroME-expressing neuron on a tiled hologram *φ*_*i*_. We then recorded, using whole-cell patch clamp recordings in organotypic slices (Supplementary Fig.6B), the photoevoked neuronal activity by *(i)* continuous illumination on *φ*_*i*_ and *(ii)* steering the laser across the *n* holograms such that each hologram is illuminated for a duration, *t*_*cyc*_, of 50 µs (Fig.4C; see Methods for details). We define two temporal parameters to characterize the two excitation conditions described above: the cell illumination time, *t*_*cell*_, as the total time during which the cell is illuminated, and the experimental time, *t*_*exp*_, as the global time needed to evoke an action potential. Under steady illumination, 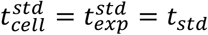, while under cyclic illumination, 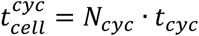 and 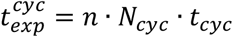, with *N*_*cyc*_the number of illumination cycles (Fig.4C).

**Figure 4:**
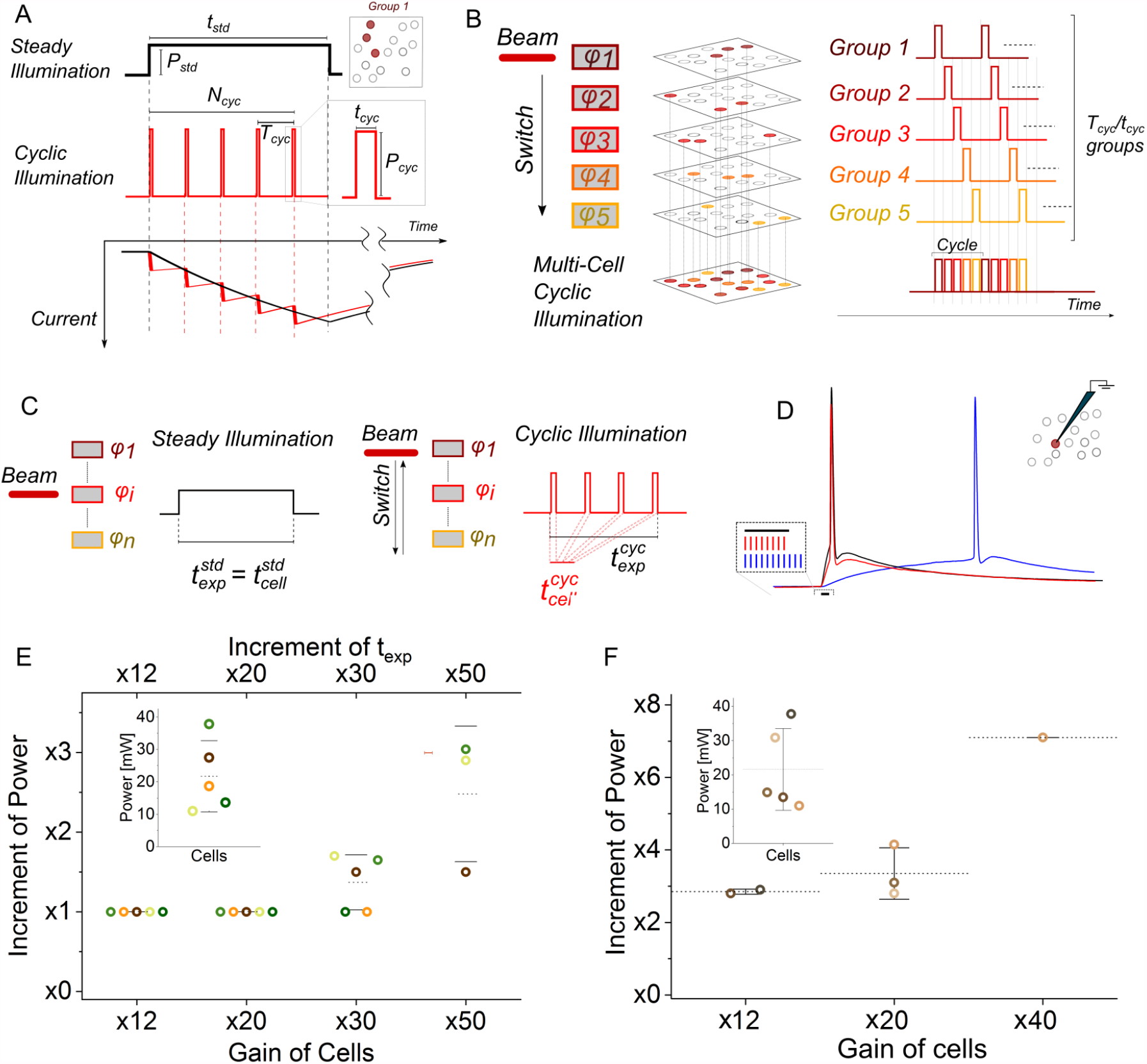
Multi-cell activation by S/P-MultiFLiT. **(A)** Photostimulation of a group of neurons under steady and cyclic illumination. A soma-targeted light pattern encoded by a single hologram can be used to photoactivate a group of neurons either under steady illumination of power *P*_*std*_ and duration *t*_*std*_ (black line, *Top*) or under cyclic illumination of power *P*_*cyc*_, period *T*_*cyc*_ and pulse duration *t*_*cyc*_ over *N*_*cyc*_ cycles (red line, *Middle*). Corresponding simulated photocurrents in an ST-ChroME-expressing neuron are shown under steady (black) and cyclic (red) illumination when 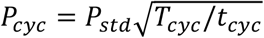 (*P*_*std*_ = 0.05 mw/µm^2^; 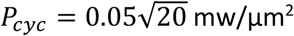; *T*_*cyc*_ = 20 *t*_*cyc*_; *t*_*cyc*_ = 50μ*s*; 1030nm) (*Bottom*). **(B)** Conceptual scheme of simultaneous photostimulation of multiple groups of neurons under Multi-S/P scheme. The LC-SLM is tiled in multiple holograms *φ*_*i*_ (here from *φ*_*1*_ to *φ*_*5*_) each encoding for different soma-targeted light-patterns illuminating different groups of cells (here from Group 1 to Group 5). The illumination beam is switched across the holograms such that each hologram is sequentially illuminated with short pulses of light *t*_*cyc*_ and the same cyclic photoactivation process is enabled sequentially on the different light patterns. The scheme displayed is meant to represent *n* groups of spots; their number is here limited to 5 for presentation purposes only. **(C)** Scheme of the experiment. The SLM is subdivided in *n* tiled holograms, with tiled hologram *φ*_*i*_ encoding for a spot of light targeting a ST-ChroME-expressing patched neuron. The neuron is then activated either by steadily maintaining the laser beam on *φ*_*i*_ with laser power *P*_*std*_ (steady illumination) or scanning the beam over the SLM with laser power *P*_*cyc*_ (cyclic illumination). **(D)** Representative light-evoked APs under steady illumination of duration *t*_*std*_=5ms (black line) and cyclic illumination with 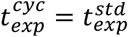 (red line) or 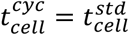 (blue line). Illumination last steadily 5ms (black bar) and cyclically 5ms (red bars) and 60ms (blue bars) **(E)** Gain of activable cells obtained in Multi-S/P for different increment of powers and 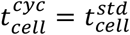. Different colors indicate different cells. Inset represents threshold power to activate the cells under steady illumination with *t*_*std*_ = 5*ms*. **(F)** Gain of number of activated cells in Multi-S/P for different increment of power and 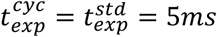. Different colors indicate different cells. Inset represents threshold power to activate the cells under steady illumination with *t*_*std*_ = 5*ms*.

At first, we optimized the excitation power, *P*_*std*_, to generate reliable APs under steady illumination for a given *t*_*std*_ = 5 ms. We found that for *P*_*std*_ = 20.4 ± 9.4 mW, we could generate APs with 7.7 ± 1.1 ms latency and 0.36 ± 0.30 ms jitter (n = 8 cells, data not shown). Secondly, we compared those values with the power, *P*_*cyc*_, and the number of cycles, *N*_*cyc*_, necessary to evoke an AP in the same cell under Multi-S/P FliT illumination (Fig.4D) by keeping either *(i)* the same excitation power (Fig.4E) or *(ii)* the same experimental time used for steady illumination (Fig.4F), i.e. either *P*_*cyc*_ = *P*_*std*_ *or* 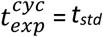, respectively.

We found that in the first condition, Multi-S/P FliT illumination can reliably generate APs in the patched cell by using up to 20 holograms (therefore in principle 20 more excitable cells) and 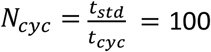, which corresponds to a total experimental illumination time 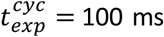 ms. Increasing the number of excitable cells up to 30 and 50 is also possible but requires increasing the excitation power *P*_*cyc*_ by a factor of ∼ 1.37 ± 0.34 and 2.48 ± 0.85, respectively (Fig.4E).

As a drawback for using 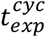 *n* times longer than *t*_*std*_, we measured large increases of AP latency and jitter (Supplementary Fig.13A). However, using 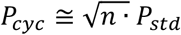 enabled keeping 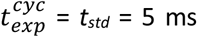 and achieving the same spiking properties as under steady illumination (Fig.4F and Supplementary Fig.13B) in agreement with the theoretical prediction (see Supplementary Note 1).

Overall, the achieved results indicate that Multi-S/P-FLiT potentially enables increasing by *n* = 20 the number of achievable cells with no increase in illumination power with respect to that used for single cell stimulation. Maintaining ms latency and sub-ms jittering is possible by using an excitation power only 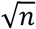 times higher than the one used for single cell excitation, which would have otherwise required *n* times higher power for conventional parallel illumination approaches.

Importantly, Multi-S/P FliT can be adapted to other cyclic illumination configurations. For instance, photostimulation protocols using low frequency, 1/ *T*_*P*_, (10-30 Hz) photostimulation train composed of short (5-10 ms) illumination pulses, *t*_*p*_ ^15,25,26^, can be equally performed using FLiT and sequential illumination of *T*_*P*_ / *t*_*P*_ tiled holograms, each for a time *t*_*P*_ (in the approximation of a switching time t ≪ *t*_*P*_). This will enable to increase by *T*_*P*_ / *t*_*P*_ the number of excited cells without incrementing the illumination power or the illumination period *T*_*P*_ (Supplementary Fig.14).

### Rise of temperature under light-driven neuronal control with FLiT

Here we demonstrated another important property of FliT illumination: the capability to minimize the light induced temperature rise for multi-target illumination.

To this end, we simulated the temperature rise under different illumination conditions using a previously validated heat diffusion model^22,23^. Firstly, we used the model to predict the temperature changes produced by 100 spots randomly distributed in a volume of 200 × 200 × 500 µm^3^ (Fig.5A) under typical illumination conditions for *in vivo* 2P optogenetics, i.e., *P*_*std*_ = 20 mW per cell and *t*_*std*_ = 5 ms. To minimize thermal crosstalk, we generated the 100 spots at an average position that enabled to maximize their relative distance. This, for a 200 × 200 × 500 µm^3^ FoE, corresponded to 50.7 ± 6.8 µm (Supplementary Fig.15A). The predicted temperature rise on the hottest spot was ∼ 3 times higher than the case of an isolated target (Fig.5C, Fig.5E and Supplementary Movie 2).

Next, we compared these findings with the case where the same 100 targets were illuminated through the sequential generation of *n* subsets of spots by keeping the same excitation conditions (i.e., *P*_*std*_ = 20 mW per target and *t*_*std*_ = 5 ms per subset). We considered two cases with sequential illumination of *n* = 4 and *n* = 10 holograms, each encoding for 25 or 10 spots, respectively (Fig.5B). Reducing the number of spots per hologram enables to further increase their average distance to 78.7 ± 14.6 µm and 106.0 ± 23.7 μm (Supplementary Fig.15B-C), and thus reducing of nearly 8% and 30% the corresponding maximum temperature rise in the hottest spot (Fig.5C, Fig.5F-G, Supplementary Fig.16 and Supplementary Movie 2). Delaying the sequential illumination by few milliseconds did not present significant variations in temperature rise compared to the previous conditions (Supplementary Fig.17). Whilst here, we have shown how to generate 100 spots in a 200 × 200 × 500 µm^3^, the concept can be extended to arbitrarily larger FoE, and correspondingly larger numbers of spots, provided that the average distance is maintained.

**Figure 5:**
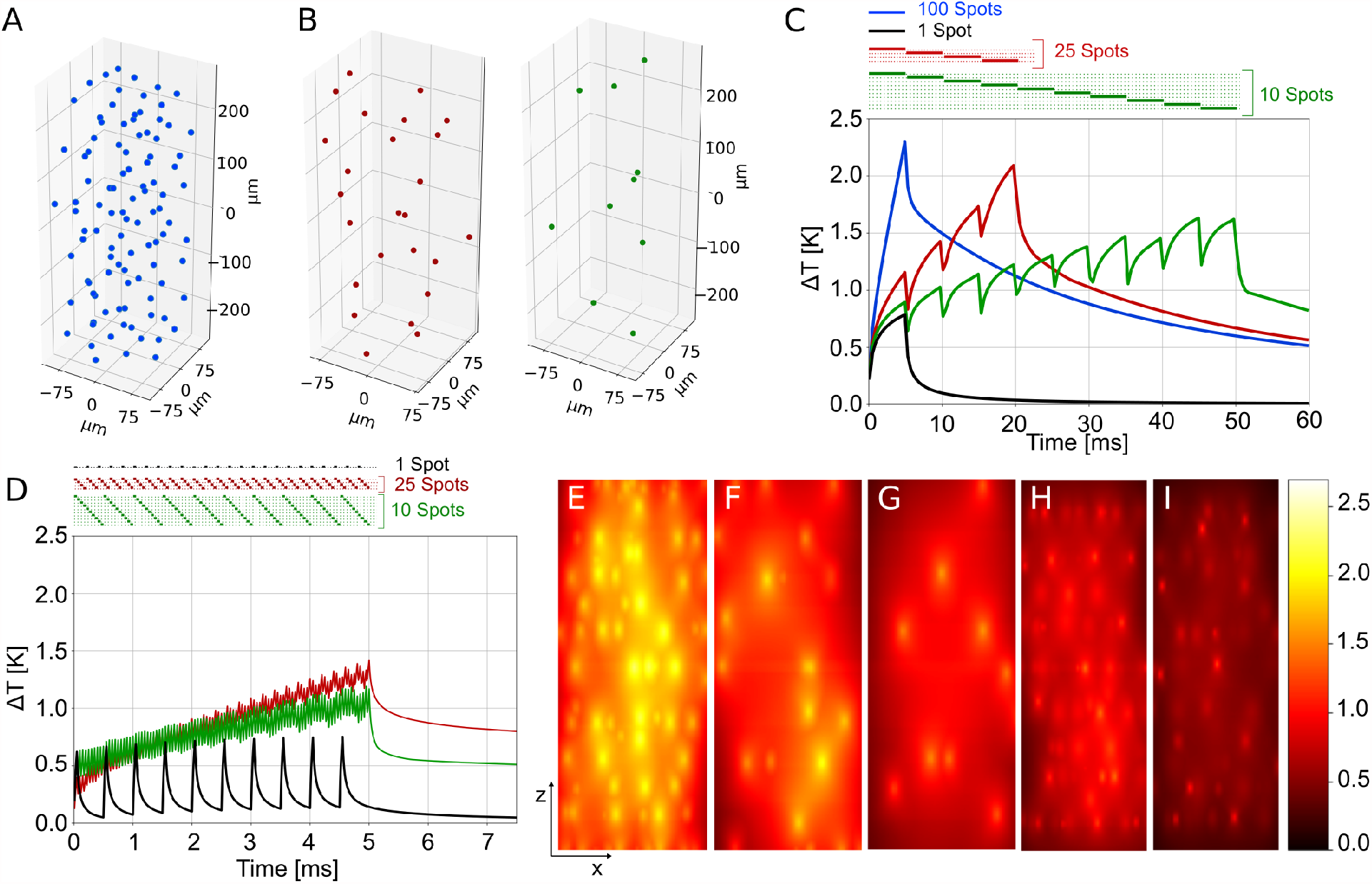
Simulated Temperature rise induced by FLiT activation under different S/P illumination protocols. **(A)** Three-dimensional view of a group of 100 spots randomly distributed in a 200×200×500µm^3^ volume. Spots were distributed to maximize nearest neighboring distance (average distance between spots equals to 50.7 ± 6.8 µm) **(B)** Representative three-dimensional view of one subset of spots when the ensemble of spots in (A) are subdivided in n=4 subsets with 25 spots each (left) or in n=10 subsets with 10 spots each (right). Average distance between spots equal to 78.7 ± 14.6 µm and 106.0 ± 23.7 for n=4 and 10, respectively. **(C)** Temperature rise in the hottest location at any given time induced by steadily illuminating 100 spots as shown in (A) either in parallel with *t*_*exp*_ = *t*_*std*_=5ms and global power *P* = 100 · *P*_*std*_ (with *P*_*std*_ =20mW) (blue) or sequentially with n=4 subsets of spots with *t*_*exp*_=20ms and global power per subset *P* = 25 · *P*_*std*_ (red) or n=10 subsets of spots with *t*_*exp*_=50ms and global power per subset *P* = 10 · *P*_*std*_ (green). The temperature rise induced by steadily illuminating one spot individually is also shown (black). The illumination timing is represented at the top of the graph with horizontal bars: different horizontal lines correspond to the timing of illumination of different subsets (illumination of one set of 100 spots (blue bar), 4 subsets of 25 spots (red bars), 10 subsets of 10 spots (green bars) and one single spot (black bar)). **(D)** Temperature rise in the hottest location at any given time induced by cyclically illuminating the 100 spots with n=4 subsets of spots for a *t*_*exp*_=5ms (*t*_*cyc*_=50µs; *N*_*cyc*_=25) and global power per subset 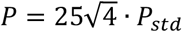 red line) or n=10 subsets of spots for a *t*_*exp*_=5ms (*t*_*cyc*_=50µs; *N*_*cyc*_=10) and global power per subset 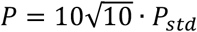 (green line). The temperature rise induced by illuminating one spot individually under cyclic illumination with the same conditions of n=10 subsets of spots is also shown (black line). The illumination timing is represented at the top of the graph with horizontal bars. Different horizontal lines correspond to the timing of illumination of different subsets (illumination of 4 subsets of 25 spots (red bars) and 10 subsets of 10 spots (green bars)). **(E-I)** xz projection of the max temperature rise produced by the 100 spots after 5 ms and simultaneous illumination of 100 spots (E); after 20 ms and sequential steady illumination with n=4 (F); after 50 ms and sequential steady illumination with n=10 (G); after 4.6 ms and cyclic illumination with n=4 (H); after 4.9 ms and cyclic illumination with n=10 (I). Color bar ranges from 0 K to 2.5 K. FoV per image 200 × 500 µm^2^.

Notably, here we have chosen a relative short illumination time. Longer illumination times (10-30 ms) will considerably lengthen the thermal diffusion length and the maximum temperature rise so that the gain in using FliT will be even more evident.

Similar reduction on the temperature rise, with no elongation of the total experimental time (*t*_*exp*_ = *t*_*std*_ = 5*ms*), can be reached if the same holograms are illuminated with cyclic illumination by using Multi-S/P FliT. In this case, we increased the excitation power per spot to 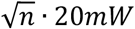 and used *t*_*cyc*_ = 50 μ*s*. Notably, cyclic illumination reduces the average temperature rises of nearly 40% and 50% for n=4 and n=10, respectively (Fig.5D, Fig.5H-I, Supplementary Movie 3 and Supplementary Fig.16) and also minimizes the temperature rise for single spot excitation.

These results show that using hybrid serial parallel photostimulation strategies (S/P FliT or Multi S/P FliT) for multi target illumination enable minimize temperature crosstalk among the multi-targets reaching a local temperature rise comparable to the case of an isolated target.

## DISCUSSION

Optical control of multiple neurons requires holographic light multiplexing through the use of LC-SLMs either coupled with spiral scanning or with parallel illumination^3,17^. In these configurations, the temporal resolution for sequential light patterning is limited by the LC refresh rate (60-500 Hz) to 2-20 ms. Moreover, although optical generation of a single neuronal spike using powers close to opsin saturation can be reached with illumination times ≤1 ms^11,24,27,28^, reaching optimal axial resolution requires working far from saturation^25^ which typically lengthens the illumination time to 5-30 ms^3^, for single spike generation, or a few seconds for the generation of multiple spikes^11^ or for neuronal inhibition^10,29^. These time values impose an extra temporal delay for sequential patterned illumination.

Holographic stimulation of multiple targets divides the laser output of high powerful lasers^21^ among multiple targets which are simultaneously illuminated. This requires taking into account possible thermal photodamages^22^ when designing the multi-site distribution. All in all, these limitations have so far restricted the maximum achievable number of targets for multi-targets 2P-optogentics to a few dozen^10,11,28^.

Here, we have presented FliT, a new scheme for multi-target excitation which overcomes all these limitations by enabling kHz projection of multiple patterns and 20 (or more) times higher number of achievable targets with respect to previously proposed holographic approaches. We have demonstrated FliT illumination in two configurations S/P-FliT and Multi-S/P-FliT where the galvanometric mirror is moved across multiple vertically aligned holograms in custom made discrete time intervals or at continuous speed, respectively.

We have shown that S/P-FliT enables to control the relative spiking time among multiple cells (or groups of cells) with a temporal delay as short as 90 μs, independently of the cell illumination dwell-time, opening the way to the investigation of synaptic integration, connectivity and neuronal coding with an unprecedented temporal precision. The ability to fast switch between multiple photostimulation patterns with sub-millisecond resolution will enable precise investigation of spatial-and time-dependent synaptic summation and integration of multiple and complex synaptic inputs^30^. Being able to stimulate multiple specific subsets of neurons, with single cell precision, either simultaneously or with sub-ms custom temporal delays will be essential to precisely probe mechanisms such as spike-time dependent plasticity (STDP), where the temporal interval between pre-and-postsynaptic spikes are necessary to strengthen or depress synaptic connections^31–34^. For instance, S/P-FliT could be used to induce STDP in adjacent spines with sub-millisecond time intervals and investigate finely the role of such processes^35^. Notably, STDP plays an important role in building specific spatiotemporal patterns involved in temporal processing, and it has been shown to be the basis for learning and memory and is known to be involved in brain pathologies^36,37^.

Previous studies on mammalian neocortex have shown that optogenetic manipulation of small (≤30 cells) groups of neurons appears sufficient to impact behavioural responses^11,12^ and most importantly that this can depend on the relative degree of synchronicity among the optically evoked spikes^16,38^. S/P-FliT has the potential to refine this type of studies by mimicking with unprecedent fine temporal precision a variety of physiological firing patterns and to manipulate them with different flavors, synchronizing or de-synchronizing them at will, while observing the effect of this time-controlled manipulation at different levels, from the local response of a neuronal circuit to behavioral responses and sensory perception, in both healthy and pathological brains.

Additionally, other brain regions with sparser connectivity and activation schemes might require the control of larger neuronal ensembles. For these studies, Multi-S/P-FliT, which enables to increase many folds the number of achievable targets, could be a crucial advance.

We have shown that Multi-S/P-FliT enables increasing *n* times the achievable number of targets, using 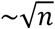 times less power than with conventional parallel illumination. This has two main implications: the possibility of using low power lasers and, for high energy laser, to reduce thermal photodamages as detailed below.

*In vivo* two photon optogenetics stimulation using mode locked Ti:Sapphire lasers (80 MHz) requires 30-50 mW/cell^29^. Considering that at the wavelengths typically used for photostimulation (i.e., 900-950 nm) these sources can provide an output of a few W (∼200 mW after the objective), multi-target stimulation using those lasers remains limited to a few cells. Multi-S/P Flit thus re-opens the possibility of using conventional mode-locked lasers to reach several dozens of spots. It also enables excitation of blue shifted opsins (PsChR2^38^, TsChR2^39^, CoChR^39^) at their optimal photostimulation peak and to combine multi-target photostimulation of these opsins with red Ca^2+^ imaging, which drastically reduces optical crosstalk from the imaging laser^13,29^.

*In vivo* activation of a single cell with spiral or parallel activation using low repetition (500 kHz-2 MHz) fiber lasers (∼1030-1064 nm excitation wavelength) requires 2-50 mW/cell^11,16,40,41^. Considering that these lasers can deliver up to 60 W, it theoretically enables simultaneous stimulation of hundreds of cells. It must however be taken into account that minimizing thermal damages^22,23^ requires reducing the thermal crosstalk among the multi targets and imposes a minimal inter-spot distance (equal to the thermal diffusion length, 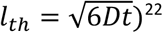)^22^. In cortical mice brain, this has so far limited to 50 cells the number of achievable targets within a 500 × 500 × 200 μm^3^ excitation volume^10^. We have shown that this limit can be overcome by decomposing the multi-target distribution into *n* sub-groups of sparser targets (i.e., average distance between targets ≫ *l*_*th*_) sequentially or cyclically illuminated via S/P Flit or Multi S/P Flit. Sequential illumination via S/P Flit enables using the same excitation power per spot, P_std_, as for the case of simultaneous illumination of the *n* sub groups but requires a *n* fold increase of the total experimental time. Cyclic illumination via Multi S/P requires increasing P_std_ by 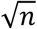 times while keeping the same total experimental time. In both cases the maximum temperature rise achieved is significantly reduced.

We have demonstrated the use of FliT for fast multi-cell optical stimulation. A similar approach can also be used for fast imaging approaches. Cohen et al.^42^ have used a gaussian beam focused with a cylindrical lens on an LC-SLM addressed with multiple tiled holograms each encoding for a specific x,y position thus achieving 2D ultrafast scanning of a diffraction limited spot. The FliT approach will enable the generation of single or multiple shaped temporally focused spots for fast multi-target imaging using e.g. voltage indicators, or for fast compressive multiphoton imaging^43^. Also, the possibility to rapidly switch between different holograms each introducing different defocusing effects can be exploited for fast repositioning of the imaging focus and ultrafast fast volumetric imaging^5,44^.

We have shown that we can tile the LC-SLM with up to 20 independent tiled holograms without a deterioration of spot quality or axial resolution. This number can be increased by including a de-scanning unit, so that each scanned hologram is projected at the centre of the objective back aperture independently of its position on the LC-SLM. This will enable to eliminate the axial tilt and intensity losses for spots generated with distal holograms. Using a lens with a shorter focal length before the LC-SLM will reduce the vertical dimension of the chirped lines on the LC-SLM and limit the losses in intensity and ellipticity observed when using n ≥20 holograms. This, eventually combined with LC-SLMs with larger pixel numbers, will enable to combine the fine temporal resolution of S/P FliT with the multi-target capability of Multi-FliT to control at fine time scale large neuronal ensembles.

In the present design, the incoming illumination has been shaped in the form of a gaussian beam. Alternatively holographic light shaping could also be used^45^ with the advantage of generating spots of variable size and shape. However, in this case, a full-frame illumination is expected in the Fourier plane of the grating (i.e., on the multiplexing LC-SLM). Hence it would be necessary to introduce an additional asymmetric focusing unit (e.g., a cylindrical lens) in order to produce tiled illuminations.

The switching unit here adopted relies on a GM. Different types of scan unit, such as polygonal scanners or AODs could be incorporated to further improve the speed of the switch between light patterns.

In conclusion FLiT illumination is a new tool for the investigation of neuronal circuits with a sub-millisecond control, at single or large neuronal population scales. Combining all the aspects of FLiT presented here, together with the latest engineered fast activity sensors, will allow an all-optical interrogation and manipulation of brain activity to decipher how specific spatio-temporal patterns produced on user defined neuronal ensembles influence specific behaviors, cognitive tasks or defined pathological conditions.

## Acknowledgments

We thank Florence Bui and Valeria Zampini for helping with stereotaxic injections and Imane Bendiffallah with organotypic cultures. We thank Dimitrii Tanese for helpful discussion on the modeling of opsin photocycles. We thank Ruth Sims for proof reading of the manuscript, Vincent de Sars for helpful discussion for the implementation of holographic algorithm in Python, Hillel Adesnik for providing the opsin ST-ChroME. We thank the IHU FOReSIGHT grant (Grant P-ALLOP3-IHU-000), the Fondation Bettencourt Schueller (Prix Coups d’élan pour la recherche française), the Getty Lab, the National Institute of Health (Grant NIH 1UF1NS107574 - 01), the Axa research funding and ERC advanced Grant HOLOVIS for financial support.

## MATERIALS AND METHODS

### Optical Setup

The optical system was built around a commercial upright microscope (Olympus BX51WI) placed on a XY stage for sample displacement (Luigs & Neumann, V380FM). A femtosecond pulsed beam delivered by a diode pumped, fiber amplifier system (Amplitude Systèmes, Goji HP; pulse width 150 fs, tunable repetition rate 10–40 MHz, maximum pulse energy 0.5 µJ, maximum average power 5 W, wavelength *λ* = 1030 nm) operated at 10 MHz, was sent first through a λ/2 wave retarder (Thorlabs, 690-1200 nm, AQWP05M-980) in combination with a polarizer cube (CVI Melles Griot) for a manual control of the laser power. The beam was then demagnified with a telescope (f1 = 100 mm; AC508-100-B, Thorlabs; f2 = 50 mm, AC508-50-B, Thorlabs) and sent through an acousto-optic modulator (AOM) (Opto-Electronic, France) to drive fast and precise light power control. The first diffracted order was projected on a pair of XY GMs (3 mm aperture, 6215H series; Cambridge Technology) with a de-magnifying telescope (M=0.4 magnification). Only the Y GM was used and driven by a servo driver (Cambridge Technology, MicroMax series 671). The GM plan was conjugated to a reflective dispersion grating of 800 l/mm by means of a telescope (f = 250 mm; AC508-250-B, Thorlabs; f = 500 mm, AC508-500-B, Thorlabs). A lens (f = 500 mm, Thorlabs, AC508-500-B) transmitted the resulting spatially chirped beam on the sensitive area of a reconfigurable liquid-crystalon-silicon LC-SLM (LCOS-SLM X10468-07, Hamamatsu Photonics, resolution 800×600 pixels, 20 μm pixel size), located in the Fourier plane of the diffraction grating. The LC-SLM was finally conjugated to the back focal plane of the microscope objective (Olympus LUMPlanFL 60XW NA 0.9) via a telescope (f = 1000 mm; AC508-1000-B, Thorlabs; f = 500 mm, AC508-500-B, Thorlabs).

The LC-SLM was divided in *n* horizontal tiles, each independently configurable. Each tiled hologram could be encoded with different sets of 3D diffraction-limited spots enabling to multiplex the temporally focused gaussian beam in multiple targeted locations on the sample. The phase profile of each *n* zones was independently calculated with a weighted Gerchberg and Saxton Algorithm^46^. The effect of the zero order in the sample was suppressed by introducing a cylindrical lens in front of the LC-SLM as detailed in^47^. Each tile of the LC-SLM was illuminated by deflecting the GM of a certain angle, corresponding to a precise driven voltage. A calibration was done in order to associate the beam position on the LC-SLM and the voltage to be applied on the GM.

During S/P-FLiT experiments for sub-ms desynchronization of pairs of neurons, the AOM and GMs was driven with a Digidata 1440A interface and pClamp software (Molecular Devices). In S/P-FLiT experiments for mimicking of random spike patterns and Multi-S/P-FLiT experiments, the system was controlled with a digital-analog converter board (National Instrument, USB-6259). The control of the system was fully automatized through a homemade software written in Python 3 and using the open graphic library PyQt5 which allowed automatic calculation of the tiled holograms and control of the the GM rotation and AOM attenuation.

### Optical Characterization of Two-Photon Excitation

In order to characterize system performance, 2PE holographic fluorescence patterns were collected by exciting a thin (∼1 μm) spin-coated layer of rhodamine-6G in polymethyl methacrylate 2% w/v in chloroform. Holographic patterns were projected on the sample plane through an excitation objective (Olympus LUMPlanFL 60XW NA 0.9). Images were collected by an opposite imaging objective (Olympus LUMPlanFL 60XW NA 0.7) in transmission geometry and detected by a CCD camera (pco, panda 4.2 bi). A short-pass filter rejected laser light (Chroma Technology 640DCSPXR; Semrock, Brightline Multiphoton Filter 680/sp). 3D stacks were collected by maintaining the excitation objective in a fixed position and moving the imaging objective along z direction with 1µm steps by means of a piezoelectric motor (MIPOS100, Piezosystem Jena).

Axial distribution of intensity on different spots was measured by integrating the pixel intensity across circular region of observations (ROIs) around the spots in each z plane. Each axial intensity distribution was fitted with a Lorentzian model. The intensity and axial resolution for each spot was evaluated and reported as maximum intensity and Full Width Half Maximum (FWHM) of the fitted curves, respectively. Images were analyzed with ImageJ and 3D rendering was performed by Imaris. Axial resolution of in-focus spots was measured by averaging the axial resolution of individual spots distributed in a two-dimensional 5×5 spots matrix in the field of excitation of each tiled hologram (30 µm inter-spots distance) as depicted in Supplementary Fig.4. In-focus intensity homogeneity of each FoE was measured by generating two-dimensional groups of 10 spots randomly distributed in the FoE of each tiled hologram. The axial resolution of spots distributed in a 3D volume was obtained by averaging the axial resolution of groups of 8 spots randomly distributed in a 120×120×70 µm for each tiled hologram. The same groups of spots were used to measure the 3D intensity homogeneity of the different tiled holograms.

### Characterization of the switching time between tiles of the LC-SLM

We characterized the switching time to reposition the beam on different tiles of the LC-SLM by means of a photodiode as schematized in Fig.2.

First, we measured the time needed to switch between adjacent tile *i* and tile *i*+*1* of the LC-SLM subdivided in 20 holograms. For that, we generated two distinct phase masks, *φ*_*i*_ and *φ*_*i*+*1*_, each encoding for an individual spot placed in a specific XY location of the focal plane. We positioned the photodiode (PD) in a conjugated plane of the sample and we aligned it such that the spot illuminates the center of the detector. We displayed *φ*_*i*_ on the tile *i* and we recorded the light intensity on the PD, while driving the GM servo with a single-step voltage pulse (pulse width 1s) which deflect the beam across small angles between tile *i* to tile *i*+*1*. We repeated the same procedure by displaying *φ*_*i*+*1*_ on tile *i*+*1*. From these two measurements, we obtained the averaged switching time to move between two consecutive tiled holograms in opposite directions, as the time taken for the signal to rise/fall between 3% and 97% of the maximum intensity. Of note, the position of PD was finely adjusted to maximize the photon counting when the GM was stationary positioned on tile *i* or tile *i*+*1*.

Second, we measured the minimal switching time between holograms when sequentially scanning at constant rate all holograms. We generated a hologram *φ*_*i*_ on a single tile *i* encoding for an individual spot detected by the PD as previously described. We then recorded the light intensity on the PD, while driving the GM servo with a staircase voltage pulse (pulse time interval 50µs) which deflects the beam across wide angles between tile *1* to tile *20*. From that, we measured the beam dwell-time on hologram *φ*_*i*_ during switch between hologram *φ*_*1*_ and hologram *φ*_*20*_. We repeated the same procedure for all 20 holograms. From that, we measured the beam dwell-time on each tiled hologram during whole scan of all holograms at constant switch rate. Of note, scan of all holograms would be alternatively possible by driving the GM with a single-step voltage facilitating maximum speed deflection of the beam across wide angles between tile *1* to tile *20*. While that can facilitate shorter dwell-time per hologram, it also gives variable dwell-time per holograms as central tiled holograms feature shorter illumination dwell-times compared to distal tiled holograms as mirror reaches maximum speed at the midpoint.

### Animals

All procedures involving animals were in accordance with national and European (2010/63/EU) guidelines and were approved by the authors’ institutional review boards and national authorities (French Ministry of Research, protocol ID: 02230.02). Experiments were performed on C57BL/6J male mice (Jackson lab.) reared in a 12 hr light/dark cycle with food ad libitum. All efforts were made to minimize suffering and reduce the number of animals.

### In Vivo Viral Expression

Stereotaxic injections of the fast somatic opsin ST-ChroME were performed in 3-week-old male mice. Mice were anesthetized with ketamine (80 mg/kg)–xylazine (5 mg/kg) solution and a small craniotomy (0.7 mm) was made on the skull overlying V1 cortex. Injection of 1μl of solution containing the viral vector was made with a cannula at a rate of 80-100 nl/min and 200-250 μm below the dural surface. We used a viral mixture containing the somatic opsin ST-ChroME (AAV9-hSyn-DIO-ChroME-Flag-ST-P2A-H2B-mRuby-WPRE-SV40, from the Adesnik lab, Berkeley, viral titer of 5.86×10^13^ particules/ml) and the Cre recombinase (AAV9-hSyn-Cre, from Addgene, 3.3×10^13^p/ml), diluted at a factor 10 and 100 respectively, in fresh NaCl solution. The craniotomy and the skull were then sutured and the mouse recovered from anesthesia. After 2-3 weeks, sufficient for an adequate expression of the virus, mice were used for electrophysiological experiments. ST-ChroME expression in acute cortical slices is shown in Supplementary Fig.6A.

### Preparation of Organotypic Cultures and Viral Infection

Hippocampal slices cultures were prepared from postnatal day 6-9 mice pups according to the interface culture method^48^. Briefly, hippocampi were gently detached from the brain and placed in a cold dissecting medium composed of: Gey’s Balanced Salt Solution (Sigma G9779), supplemented with 25 mM D-glucose, 10 mM HEPES, 1 mM Na-Pyruvate, 0.5 mM α-tocopherol, 20 nM ascorbic acid and 0.4% penicillin/streptomycin (5000 U/mL; Fisher 11528876). Transverse slices of 300 μm tickness were cut using a McIlwain tissue chopper, maintained for at least 1h at 4°C and then transferred onto semiporous membranes inserts (47 mm diameter, 0.45 µm pore size; Millipore FHLP04700) which were placed in six well tissue culture plates containing 1.1 ml medium per well. The incubation medium consisted in: 50% Opti-MEM (Fisher 15392402), 25 % heat-inactivated horse serum (Fisher 10368902), 24% HBSS (Fisher 15266355), 1% penicillin/streptomycin (5000U/mL), and supplemented with 25 mM D-glucose, 1 mM Na-Pyruvate, 20 nM ascorbic acid and 0.5 mM α-tocopherol. Slices were maintained at 34°C in an incubator with 5% CO_2_. After 3 days, the medium was replace with a fresh and warm Neurobasal culture medium composed of: 2% Neurobasal-A (Fisher 11570426), 15% heat-inactivated horse serum, 2% B27 supplement (Fisher 11530536), 1% penicillin/streptomycin (5000U/mL), and supplemented with 0.8 mM L-glutamine, 0.8 mM Na-Pyruvate, 10 nM ascorbic acid and 0.5 mM α-tocopherol. This medium was changed every 2-3 days until the experiment.

Organotypic slices were then infected with 1 μL of virus at 5-7 days in vitro (DIV). We used the same mixture as for in vivo stereotaxic injections. Slices were used for electrophysiology recordings at 12-14 DIV. See Supplementary Fig.6B for ST-ChroME expression in this preparation.

### Acute Slice Preparation for Electrophysiology

Acute parasagittal slices of the visual cortex were prepared from adult mice 2-3 weeks after viral injection. Animals were decapitated after being deeply anesthetized with isoflurane (5% in air). The brain was quickly removed, immersed in an ice-cold choline solution and 300 μm-thick slices were obtained using a vibratome (Leica Biosystems VT1200S). The cutting solution contained the following (in mM): 126 choline chloride, 16 glucose, 26 NaHCO_3,_ 2.5 KCl, 1.25 NaH_2_ PO_4_, 7 MgSO_4_, 0.5 CaCl_2_, pH 7.4, cooled to 4°C and equilibrated with 95% O_2_ /5% CO_2_. Slices were maintained at 32°C for 20min in standard ACSF (sACSF) containing the following (in mM): 125 NaCl, 2.5 KCl, 26 NaHCO3, 1.25 NaH2PO4, 1 MgCl2, 1.5 CaCl2, 25 glucose, and 0.5 ascorbic acid, pH 7.4, saturated with 95% O2 and 5% CO2 and then transferred at room temperature in the same solution until recordings.

### Whole-Cell Electrophysiology In Vitro

Acute slices as well as organotypic slices were placed in a recording chamber under the microscope objective, and perfused continuously with fresh sACSF saturated with 95% O2 and 5% CO2. Neurons were patched at 30-60 μm from the slice surface. Single or doubled-patched neurons were clamped at −70 mV in voltage-clamp configuration and membrane potential was kept at −70 mV with currents injections in current-clamp configuration. Patch electrodes (Borosilicate glass pipette, outer diameter 1.5 mm and inner diameter 0.86 mm, Sutter Instruments) were filled with an intracellular solution containing the following (in mM): 127 K-gluconate, 6 KCl, 10 Hepes, 1 EGTA, 2 MgCl2, 4 Mg-ATP, 0.3 Na-GTP; pH adjusted to 7.4 with KOH. The estimated reversal potential for chloride (E_Cl_) was approximately −69 mV based on the Nernst equation. Pipettes were pulled from borosilicate glass capillaries and had a typical tip resistance of 5-6 MΩ. The averaged serie resistances were 18.5 ± 7.9 MΩ (n = 34 cells) and 17.9 ± 3.7 MΩ (n = 8 cells), for acute slices and organotypic cultures, respectively. The following receptor blockers were added to the sACSF to block any synaptic effect: DNQX and AP-V (1µM each; from Abcam). Electrophysiology data were acquired with a Multiclamp 700B amplifier and digitized with a Digidata 1322A interface and pClamp software (Molecular Devices). Signals were sampled at 20–50 kHz and filtered at 4-10 kHz.

### Desynchronization of Activity of distinct neurons

In S/P-FliT experiments, we desynchronized two ST-ChroME-expressing targeted neurons, here called neuron A and neuron B (Fig.3A). The following photostimulation procedure was used in order to trigger activity in neuron A and B with a time delay shorter than the illumination dwell-times needed to evoke activity in the two neurons. We defined three tiled phase masks and we vertically piled them adjacently on the LC-SLM display such that: tile φ_A_ encodes for illumination of neuron A (top tile), tile φ_AB_ encodes for simultaneous illumination of neurons A and B (middle tile), and tile φ_B_ encodes for illumination of neurons B (bottom tile). First, we established threshold light powers P_A_ and P_B,_ and illumination dwell-times t_A,_ t_B_ to independently evoke an AP on neuron A and neuron B, by deflecting the GM on tile φ_A_ and φ_B_. Threshold values were defined in current clamp mode when AP was reliably generated on 3/3 consecutive trials (40 s inter-time between trials). Photocurrents corresponding to threshold illumination conditions were also recorded in voltage-clamp. On the basis of these values, we set a sequence to drive the GM and the AOM and introduce arbitrarily defined spike delays δt between neuron A and B. Accordingly, the beam was sequentially directed by tilting the GM on tile φ_A_ for a time δt, on tile φ_AB_ for a time *t*_*A*_ − δt and on tile φ_B_ for a time *t*_*B*_ − (*t*_*A*_ − δt). The incoming power was adjusted via the AOM such that it was set to P_A_ when the beam was on tile φ_A_, to P_B_ when the beam was on tile φ_B_ and to *P*_*A*_ + *P*_*B*_ when the beam was on tile φ_AB_. Of note, the diffraction efficiency of phase mask φ_AB_ was computationally corrected such that the ratio of intensity sent onto neuron A and B equals to *P*_*A*_ /*P*_*B*_ That ensures that both neurons are constantly illuminated with the same intensity during their respective illumination dwell-time. Importantly, in voltage-clamp mode, we verified that the beam of power *P*_*A*_ + *P*_*B*_ positioned on φ_AB_ for a time *t*_*A*_ and *t*_*B*_ elicited the same photocurrents previously elicited by illuminating only neuron A (with *P*_*A*_ power, *t*_*A*_ dwell-time on φ_A_) and B (with *P*_*B*_ power, *t*_*B*_ dwell-time on φ_B_), respectively (Supplementary Fig.18). GM deflection between the three tiles was driven with small angle single-step voltage as previously detailed. We thus recorded in current-clamp the APs driven in neuron A and B by addressing GM and AOM following the established sequence of photoactivation.

Of note, in general, for all those experiments which feature delays longer than the illumination dwell-times needed to evoke activity in the two neurons, only two tiles of the LC-SLM are necessary (tile φ_A_ and tile φ_B_), as the beam will never be simultaneously on neuron A or B.

Importantly, this strategy can be generalized to desynchronize *n* neurons (or n groups of neurons) with delays inferior to each activation dwell-time, by dividing the LC-SLM in 2*n* − 1 tiled holograms and piling them on the LC-SLM such that each hologram encodes, from top to bottom: *1*^*st*^ tiled hologram → *1*^*st*^ neuron, *2*^*nd*^ tiled hologram → *1*^*st*^+*2*^*nd*^ neurons,…, *n*^*th*^ tiled hologram → *1*^*st*^+*2*^*nd*^+*…*+*n*^*th*^ neurons, *(N*+*1)*^*th*^ tiled hologram → *2*^*nd*^+*…*+*n*^*th*^ neurons, *(n*+*2)*^*th*^ tiled hologram → *3*^*rd*^+*…*+*n*^*th*^ neurons … *(2n-1)*^*th*^ tiled hologram → *n*^*th*^ neuron. Power on each of the 2n-1 tiled phase masks needs to be modulated accordingly to the number of encoded targets (Supplementary Fig.8).

### Mimicking of firing

In S/P FliT experiments aiming to mimic neuronal firing, reference traces originated from an individual recording under in-vivo patch-clamp. In particular, two subsections, each 2 s long and featuring characteristic firing patterns, were arbitrarily selected and delayed. The two traces were then feed to a home-made software which extracted the spike timing and automatically determined the illumination sequence (including illumination power and switch time) to be addressed on the tiled holograms of the LC-SLM to reproduce the delayed spiking patterns on two double-patched neurons.

### Multi-Neuron Activation

In Multi-S/P experiments, the LC-SLM was subdivided in *n* tiled phase masks. In particular, one mask *φ* was encoded to illuminate one targeted ST-ChroME-positive neuron. We initially established threshold light power P_std_ and illumination dwell-time t_std_ to evoke an AP on the cell by tilting the GM to steadily illuminate *φ*.

The cell was then photoactivated under cyclic illumination by driving the GM with a staircase voltage input which facilitated steering the beam back and forth on all *n* holograms through discrete angle deflections and fixed dwell-time per hologram at *t*_*cyc*_ = 50*μs* for *N*_*cyc*_ cycles (Supplementary Fig.19). Compared to scan the holograms by driving the GM with a single-step voltage input, staircase voltage inputs gives more homogenous dwell-time per hologram.

We tested a slow photoactivation protocol featuring a total scan time 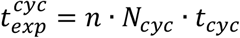 and a fast photoactivation protocol featuring a total scan time 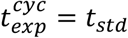. We established power to trigger an AP in both cases by varying the number of the tiled holograms (i.e., the size of each tiled hologram) between 12 and 50. Photocurrents have been recorded in voltage-clamp by displaying *φ* on different position of the LC-SLM, in order to verify that different tiles substantially elicit the same photocurrent (Supplementary Fig.11).

### Temperature Simulation

The spatio-temporal distribution of the temperature rise was calculated by solving the Fourier heat diffusion equation^49^ considering the brain tissue as an infinite medium with isotropic and uniform thermal properties as described in ^22^. The solution is obtained by convolving the Green’s function for the diffusion equation by the thermal source term, which is the thermalisation of the absorbed light source intensity. This model has been experimentally validated^22^. Numerical solution was implemented in Python, taking special care in selection spatial and temporal sampling to avoid overlap due to cyclic boundary conditions induced by the use of Fourier transform based numerical convolution.

### Data Analysis and Statistical Tests

We performed the analysis of the recorded stacks on Rhodamine layers with MATLAB, ImageJ, and the Imaris software (Bitplane, Oxford Instruments). The 2PE fluorescence values for each spot were obtained by integrating the intensity of all the pixels in a circular area containing the spot, in the plane where the intensity was at its maximum value (i.e., the TF plane). Axial intensity distributions were obtained by integrating the intensity of the pixels in the same area for each plane of the recorded stack, in a range of ± 20 μm around the focal plane of each spot. Reported values for the axial confinement were the fit of the axial profile of the spots with a Lorentzian model and referred to the FWHM of the curves.

All electrophysiological data were analyzed with Clampfit (Molecular Devices). For S/P-FliT experiments (Fig.3), we measured, for double patched neurons, A and B, the depolarization onset or the AP peak delay, determined as the time between the beginning of the light stimulus and membrane potential change or the AP peak, respectively. We then substracted the values of cell B to cell A and compared this temporal delay to the expected one. We evaluated the temporal accuracy as the difference between imposed δt and experimental 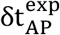 delays, 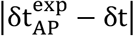. Global accuracy was calculated as weighted mean and SD of all imposed δt. For mimicking experiments, the analysis of the results was established by pairing the closest subsequent APs in the two neurons. In particular, for each AP pair, we evaluated the temporal accuracy as the difference between driven and experimental inter-spike time. We then calculated the overall accuracy of the mimicking by weight averaging the temporal accuracy of each AP pair.

AP latencies were determined as the time between the beginning of the stimulus and the time of AP peak, and AP jitters were calculated as the standard deviation (SD) of the AP latency accros trials. All recordings were analyses and averaged across 3-5 photostimulation trials. All values are presented as mean ± SD of *n* experiments.

