## Supplementary Information for "Ultrafast Light Targeting for High-Throughput Precise Control of Neuronal Networks"

**SUPPLEMENTAL INFORMATION**

### Supplementary Note 1

The dynamic of photocurrent can be described using a four-state model which assumes two open and two closed states[50–53] (Supplementary Fig.20A).

In this case, by denoting with $O_1$, $O_2$, $C_1$, and $C_2$ the fraction of opsin molecules in each of the four states at any given instant of time, the transition rates among the four states can be described by the following set of rate equations,

$$\begin{aligned} \dot{O}_1 &= G_{a1}(P)C_1 + G_b(P)O_2 - \left(G_{d1} + G_f(P)\right)O_1 \\ \dot{O}_2 &= G_{a2}(P)C_2 + G_f(P)O_1 - \left(G_{d2} + G_b(P)\right)O_2 \\ \dot{C}_1 &= G_{d1}O_1 + G_r C_2 - G_{a1}(P)C_1 \\ \dot{C}_2 &= G_{d2}O_2 - \left(G_r + G_{a2}(P)\right)C_2 \end{aligned} \quad (1)$$

where $O_1 + O_2 + C_1 + C_2 = 1$ and $G_{a1}(P)$, $G_{a2}(P)$, $G_{d1}$, $G_{d2}$, $G_f(P)$, $G_b(P)$ and $G_r$ are the rate constants for transitions $C_1 \rightarrow O_1$ , $C_2 \rightarrow O_2$ , $O_1 \rightarrow C_1$ , $O_2 \rightarrow C_2$ , $O_1 \rightarrow O_2$ , $O_2 \rightarrow O_1$ and $C_2 \rightarrow C_1$ respectively.

Under 2P excitation, the dependence on the excitation power, P, of the transition rates from closed to opened states, $G_{a1}(P)$ and $G_{a2}(P)$, can be expressed as[50]:

$$G_{a1}(P) = k_1 \frac{P^2}{P_m^2 + P^2} \ ; G_{a2}(P) = k_2 \frac{P^2}{P_m^2 + P^2} \quad (2)$$

and between the opened states as:

$$G_f(P) = k_f \frac{P^2}{P_m^2 + P^2} + G_{f0} \ ; G_b(P) = k_b \frac{P^2}{P_m^2 + P^2} + G_{b0} \quad (3)$$

Where $P_m$ marks the transition from quadratic to saturated regime for rate constants and $k_1$, $k_2$, $k_f$ and $k_b$ are the transition rate at saturation of $C_1 \rightarrow O_1$ , $C_2 \rightarrow O_2$, $O_1 \rightarrow O_2$ , $O_2 \rightarrow O_1$, respectively.

Following the procedure described in Evans et al.[50] we fitted the photocurrent traces obtained upon targeted 2PE illumination of a ST-ChRoME-expressing neuron and determined the transition rate constants for the opsin ST-ChRoME (Supplementary Fig.20B). We then used these values to simulate the photocurrents induced in a ST-ChroME-expressing neuron by using a steady or cyclic illumination for different illumination powers and number of tiled holograms *n* (Supplementary Fig.20C). We found, that for intensities P << P $_m$ and short cyclic flashes of light, $t_{cyc}$, photocurrents evoked under cyclic illumination are comparable to the ones obtained under steady illumination if $P_{cyc} \cong P_{std}\sqrt{n}$ (Supplementary Fig.20D).

Interestingly, the same condition can be analytically derived by using a simplified two-state model, which only considers the O1 and C1 states. This assumption is valid when the illumination times are short compared to the decay times constants of the state $O_1$ (i.e., $t_{cyc} << \frac{1}{G_f(P)}$ , $\frac{1}{G_{d1}}$).

In this assumption, the rate equations (1) can be simplified in:

$$\begin{aligned}\dot{O}_1 &= G_{a1}(P)C_1 - G_{d1}O_1 \\ \dot{C}_1 &= G_{d1}O_1 - G_{a1}(P)C_1\end{aligned} \tag{4}$$

For a steady illumination power $P_{std}$ and an illumination time $n \cdot t_{cyc}$ ($n$ the number of tiled holograms and $t_{cyc}$ = 50 µs) the fraction of opsins in O1 state during the first illumination cycle is given by

$$O_{1,std}\left(n \cdot t_{cyc}\right) = \frac{G_{a1}(P_{std})}{G_{a1}(P_{std})+G_{d1}} \times \left[1 - exp\left(-(G_{a1}(P_{std}) + G_{d1}) \cdot n \cdot t_{cyc}\ \right)\right] \tag{5}$$

which considering short illumination time, $(G_{a1}(P_{std}) + G_{d1}) \cdot n \cdot t_{cyc} \ll 1$, becomes:

$$O_{1,std}\left(n \cdot t_{cyc}\right) \approx G_{a1}(P_{std}) \cdot n \cdot t_{cyc} \tag{6}$$

For a cyclic illumination of power $P_{cyc}$, and an illumination time $t_{cyc}$, the opsins are excited to the state O1 during $t_{cyc}$ and decay back to C1 during $(n-1) \cdot t_{cyc}$. The fraction of opsins in O1 state is then:

$$O_{1,cyc}\left(n \cdot t_{cyc}\right) = \frac{G_{a1}(P_{cyc})}{G_{a1}(P_{cyc})+G_{d1}} \underbrace{\left[1 - exp\left(-\left(G_{a1}\left(P_{cyc}\right) + G_{d1}\right)t_{cyc}\right)\right]}_{excitation} \cdot \underbrace{exp\left(-G_{d1}(n-1)t_{cyc}\ \right)}_{decay} \tag{7}$$

which can be approximated to

$$O_{1,cyc}\left(n \cdot t_{cyc}\right) \approx G_{a1}\left(P_{cyc}\right)t_{cyc}\left(1 - G_{d1}(n-1)t_{cyc}\right) \tag{8}$$

In order to obtain the same number of opsins in the open state under steady and cyclic illumination, i.e. $O_{1,std} = O_{1,cyc}$, two conditions need to be verified: the number of transitions from C1 to O1 are the same under steady and cyclic illuminations, i.e.:

$$G_{a1}(P_{std}) \cdot n \cdot t_{cyc} = G_{a1}\left(P_{cyc}\right) \cdot t_{cyc} \tag{9}$$

and, in the cyclic configuration, the number of transitions from O1 to C1 in the off-time $(n-1)t_{cyc}$, is negligeable:

$$G_{d1}(n-1)t_{cyc} \ll 1 \tag{10}$$

In the limit $P \ll P_m$, $G_{a1}(P)$ (equation (2)) can be approximated by:

$$G_{a1}(P) = k_1 \frac{P^2}{P_m^2 + P^2} \approx \frac{k_1}{P_m^2} P^2 \tag{11}$$

and Equation (9) can be written as:

$$\frac{k_1}{P_m^2} P_{std}^2 \cdot n \cdot t_{cyc} = \frac{k_1}{P_m^2} P_{cyc}^2 \cdot t_{cyc} \tag{12}$$

Which gives the relationship between cyclic and steady power as numerically derived using the four-states model,

$$P_{cyc} = P_{std}\sqrt{n} \tag{13}$$

**SUPPLEMENTARY FIGURES:**

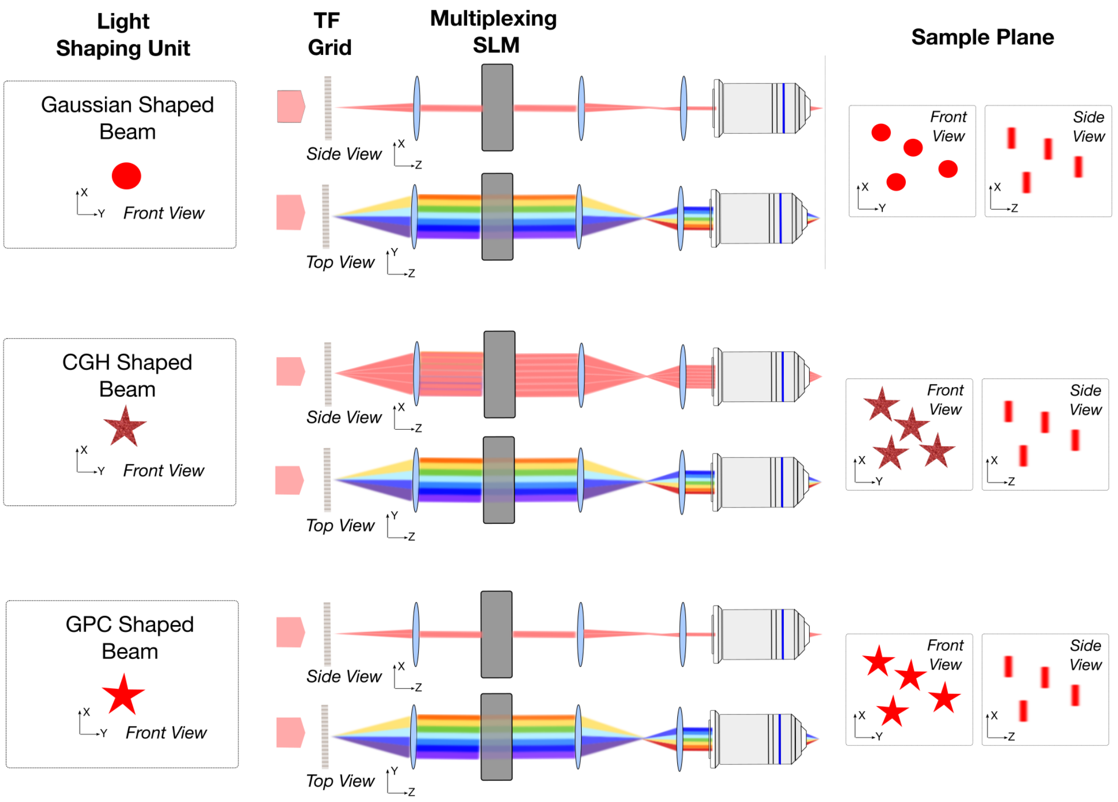


**Supplementary Figure 1:**

**Multiplexing temporally focused light patterns by using different light shaping units.**

Gaussian- (*top*), Computer Generated Holography, CGH- (*middle*) and Generalized Phase Contrast, GPC- (*bottom*) based multiplexing temporally focused systems. For each beam shape, it is here shown: XY *front* view of the incoming beam on the TF grating plane (*left*), XZ *side* and YZ *top* view of the multiplexing pathway (*center*), XY *front* and XZ *side* views of the XYZ multiplexed patterns on the sample plane (*right*).

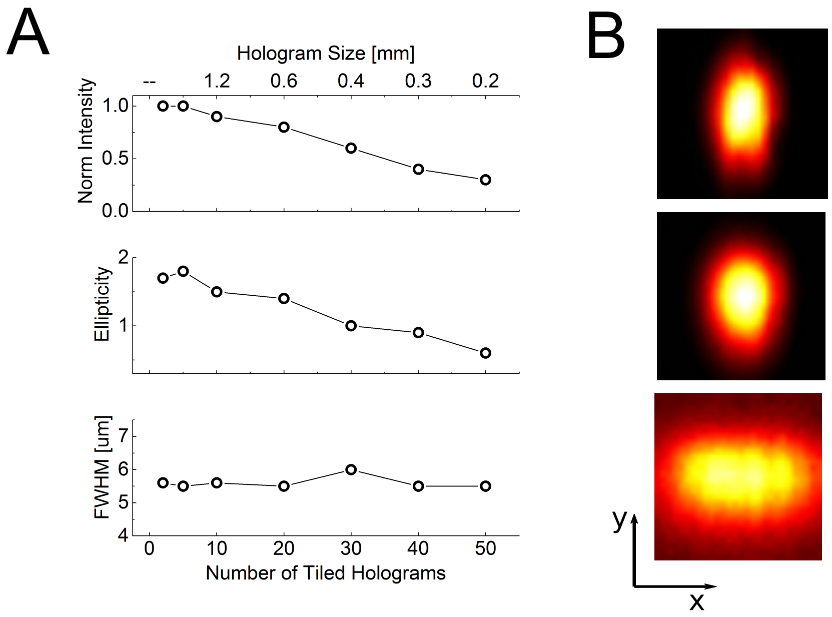


**Supplementary Figure 2:**

**Intensity, in-focus XY ellipticity and axial FWHM of the multiplexed spot vs size of tiled holograms on the LC-SLM.**

**(A)** Intensity (*Top*), Ellipticity (*Middle*) and FWHM of axial intensity distribution (*Bottom*) of a spot encoded by holograms of different sizes in the direction orthogonal to the grating dispersion. **(B)** xy images of the spot encoded by using 60pixelsx800pixels (i.e., 1.2mm x 12mm) holograms (corresponding to LC-SLM tiled in 10 regions) (*Top*), 30x800 pixels (i.e., 0.6mm x 12mm) holograms (corresponding to LC-SLM tiled in 20 regions) (*Middle*) and 12x600 pixels (0.24mm x 12mm) holograms (corresponding to LC-SLM tiled in 50 regions) (*Bottom*). LC-SLM pixel size 20µm.

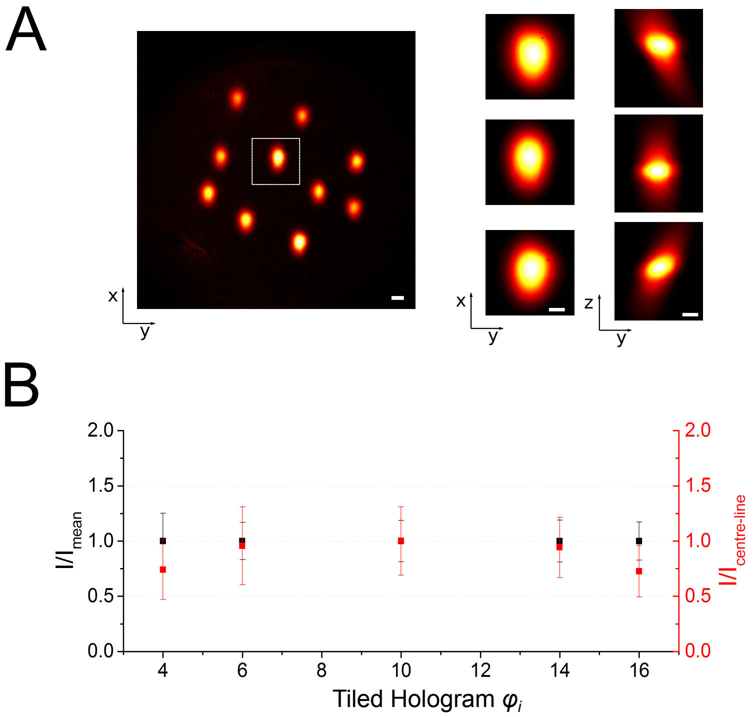


**Supplementary Figure 3:**

**Intensity distribution of groups of spots randomly distributed in the field of excitation generated by tiled holograms at different vertical position of the LC-SLM**.

(**A**) *(Left)* Representative distribution of multiple spots encoded with the hologram $\varphi_{10}$. Scale bar 10µm. *(Right)* xy and xz projections of the central spot encoded by $\varphi_4$ (Top), $\varphi_{10}$ (Middle) and $\varphi_{16}$ (Bottom). Scale Bar 5µm. **(B)** Intensity distribution of 10 random spots arranged as in (A) within (black axis) and between (red axis) the field of excitation (FoE) of different holograms $\varphi_i$ encoded on different tiles *i* of the LC-SLM. Black symbols indicate per each hologram $\varphi_i$, normalized Mean ± SD of the ratios between the intensity *I* of each spot and the averaged spot intensity $I_{mean}$ within the FoE of $\varphi_i$. Red symbols indicate per each hologram $\varphi_i$, normalized Mean ± SD of the ratios between the intensity of each spot and the intensity of the same spot encoded by the central hologram $\varphi_{10}$. Each hologram was 30x800 pixels, i.e. the LC-SLM was tiled into 20 regions.

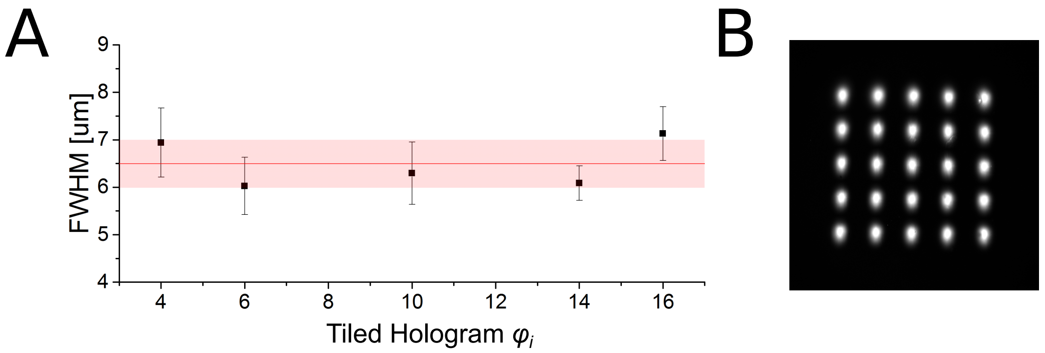


**Supplementary Figure 4:**

**Axial FWHM of multiplexed spots generated by different tiled holograms of the LC-SLM**

**(A)** Axial intensity FWHM of individual spots distributed in a matrix as in (B) and generated by different holograms $\varphi_i$. Black symbols indicate Mean ± SD of FWHM per each holograms $\varphi_i$. Red line and reddish band indicates the global mean and SD over the different $\varphi_i$, respectively. **(B)** Representative distribution of spots generated by hologram $\varphi_{10}$.

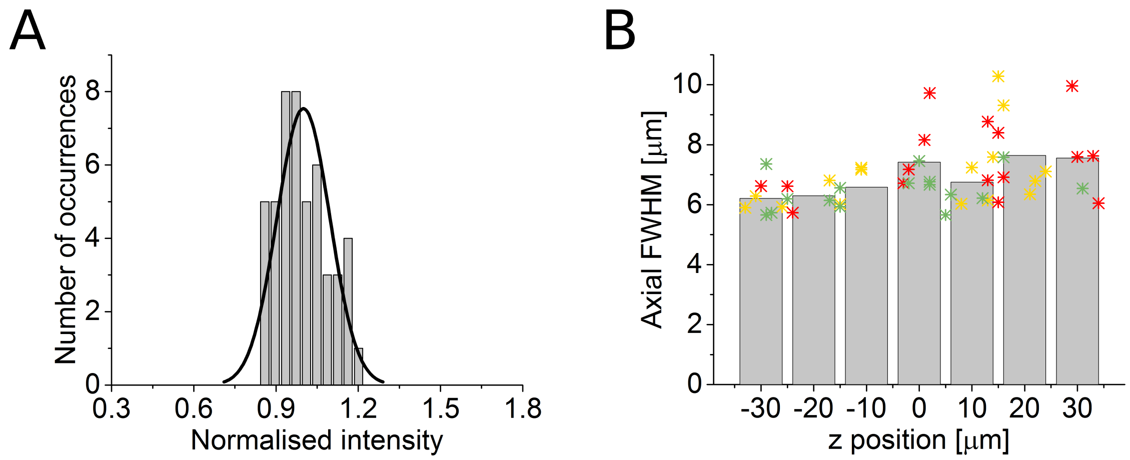


**Supplementary Figure 5:**

**Intensity distribution and axial FWHM of multiplexed spots generated by different tiled holograms of the SLM in a 3D volume.**

**(A)** Histogram of the maximal 2PE fluorescence intensity for each spot, normalized to the average intensity of all spots, after diffraction efficiency correction (48 spots randomly distributed in a 120x120x70 $\mu m^3$). **(B)** Axial confinement, calculated as the FWHM of the axial intensity profile of each spot (48 spots randomly distributed in 120x120x70 $\mu m^3$). Stars indicate FWHM of each spot. Different colors indicate spots encoded by different tiled holograms $\varphi_i$ (hologram $\varphi_4$, yellow; hologram $\varphi_{10}$, red; and hologram $\varphi_{16}$, green). Grey bars indicate the mean values in 10 µm range around the designated z position. SLM subdivided in 20 tiled holograms.

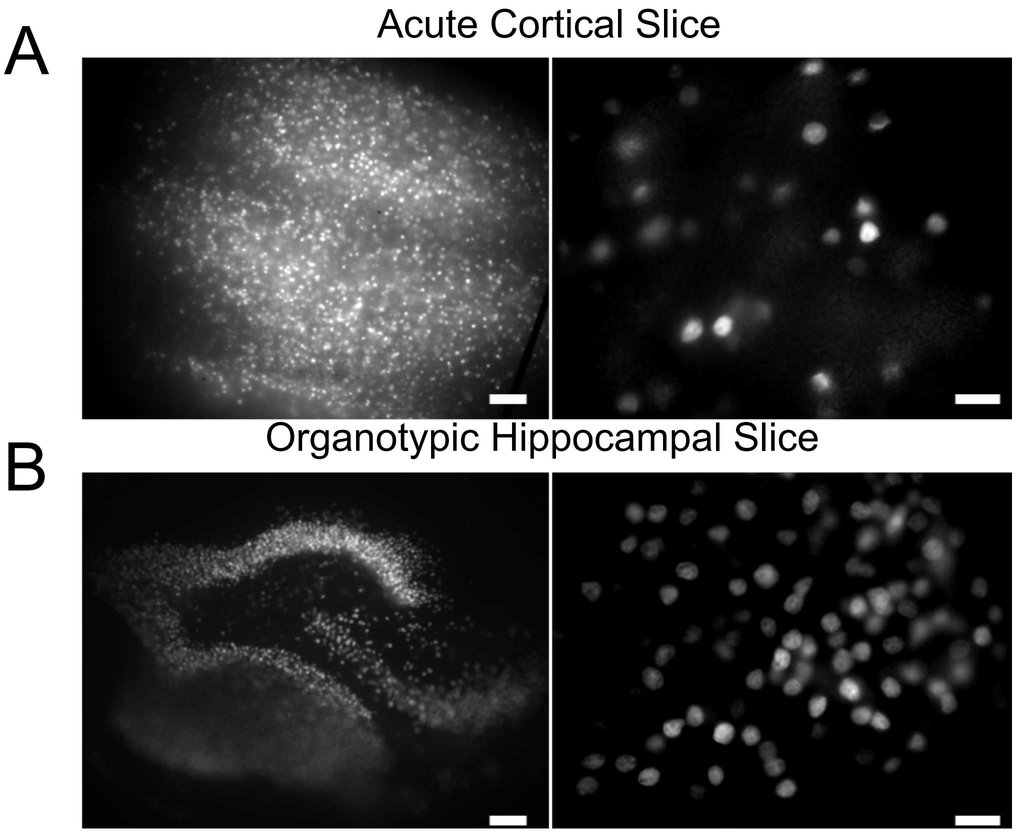


**Supplementary Figure 6:**

**ST-ChroME expression.**

**(A)** Widefield fluorescence images of ST-ChroME-positive cells in an acute cortical slice and **(B)** in an organotypic hippocampal slice. 300µm-thick slices. Left panels: scale bar 100µm. Right panels: scale bar 20µm.

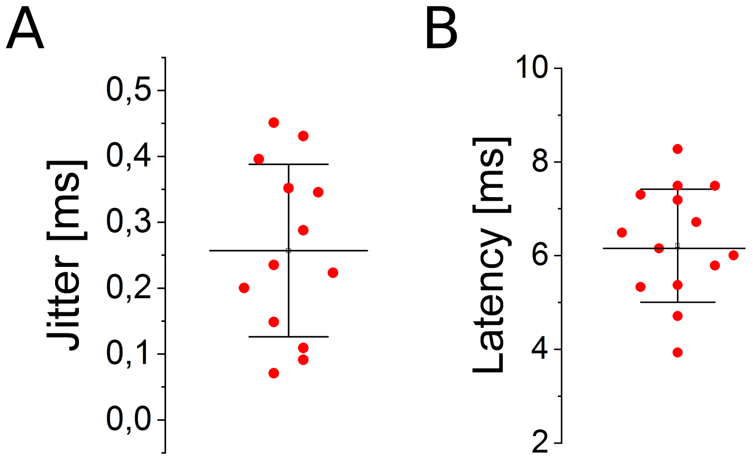


**Supplementary Figure 7:**

**Action potential jitter and latency upon soma-targeted 2PE photostimulation**.

**(A)** AP jitter and **(B)** latency for different ST-ChroME-expressing patched cells in acute cortical slices illuminated for 4-5ms dwell-time with a soma-targeted spot. Mean jitter is 0.25 ± 0.13 ms and mean latency is 6.2 ± 1.2 ms. Mean power 30.5 ± 13.6 mW. Different circles indicate different cells (n = 13). Data are shown as mean ± SD. 1030nm illumination has been used.

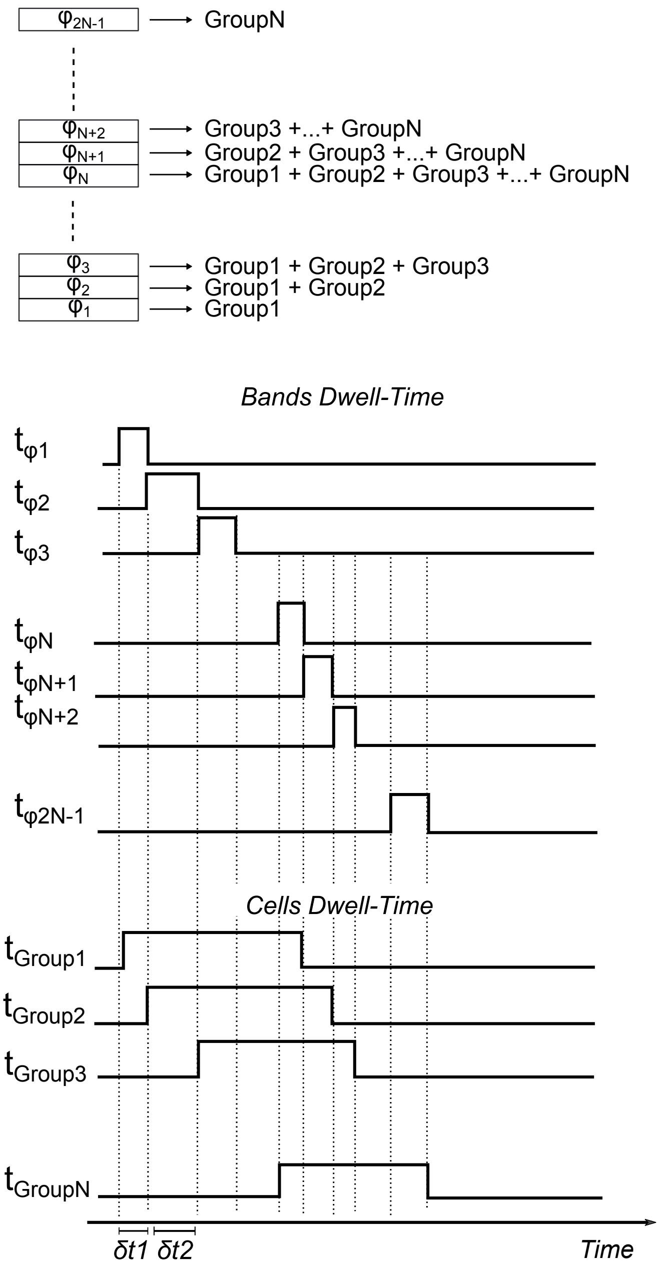


**Supplementary Figure 8: FLiT configuration to desynchronize *n* groups of neurons with delays inferior to each activation dwell-time**

**(A)** Scheme of the disposition of the tiled holograms $\varphi_i$ enabling to desynchronize *n* groups of neurons with delays $\delta t_i$ inferior to each activation dwell time. The LC-SLM is divided in $2n - 1$ tiled holograms, each encoding to target different pools of neurons, such that the *n* different groups of neurons are photostimulated individually or in parallel on the basis of their activation chronological order as depicted in (B). **(B)** Time flow of the activation of the *n* different groups of neurons. Of note, power needs to be adapted on each tiled hologram such that each groups of neurons are constantly illuminated during their activation interval.

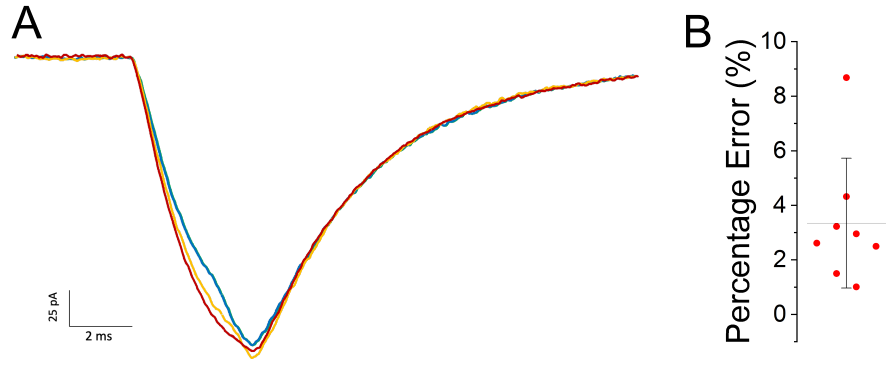


**Supplementary Figure 9:**

**Light-induced currents during switch between the different holograms.**

**(A)** Representative photocurrents from a ST-ChroME-expressing neuron when illuminated for 6ms with a single spot encoded by a single tiled hologram $\varphi_i$ (red line) or by switching between three adjacent tiled holograms, each encoding the same spot, with different dwell time per holograms (blue line: $\varphi_i$=2ms + $\varphi_{i+1}$=2ms + $\varphi_{i+2}$=2ms; green line: $\varphi_i$=1ms + $\varphi_{i+1}$=4ms + $\varphi_{i+2}$=1ms; yellow line: $\varphi_i$=3ms + $\varphi_{i+1}$=2ms + $\varphi_{i+2}$=1ms). The SLM was here subdivided in 20 tiled holograms. **(B)** Amplitude percentage error between photocurrents induced using $\varphi_i$ for 6ms (red in panel A) or by switching between the 3 different tiled holograms each for 2ms (i.e., $\varphi_i$=2ms, $\varphi_{i+1}$=2ms, $\varphi_{i+2}$=2ms, blue in panel A). Different circles represent different cells. Mean 3.35 ± 2.38 %. n = 8 cells. Incoming illumination power was maintained constant during the switches.

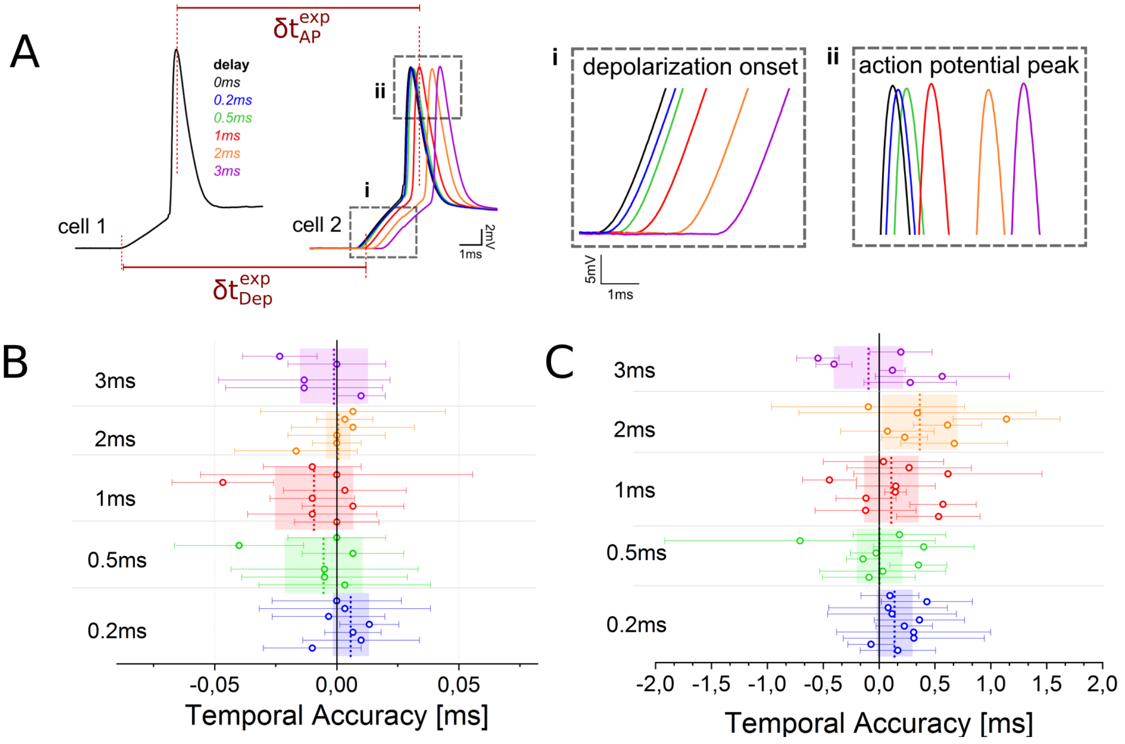


**Supplementary Figure 10:**

**Temporal accuracy of FLiT driven delays.**

**(A)** Representative traces from two ST-ChroME-expressing patched neurons illuminated with soma-targeted beams with delays δt between 0.2 and 3ms. Experimental AP peak $\delta t_{AP}^{exp}$ and depolarization onset $\delta t_{Dep}^{exp}$ delays are highlighted. **(B-C)** Temporal accuracy of depolarization (B) and AP occurrences (C) calculated for different couple of cells as $\left|\delta t_{AP}^{exp} - \delta t\right|$, and $\left|\delta t_{Dep}^{exp} - \delta t\right|$, respectively. Different circles correspond to different pairs of cells. Data are shown as mean $\pm$ SD. Different colors correspond to different delays. Vertical dashed lines and bands indicate average and SD temporal accuracy of all pairs of cells activated with the same delay time. Global mean depolarization accuracy is 1.4 ± 5.1 µs and mean AP accuracy is 96 ± 114 µs (n = 12 pair of cells). Mean photostimulation power is 36.8 ± 20.9 mW. Illumination dwell-time ranges between and 4-5ms. 1030nm illumination has been used.

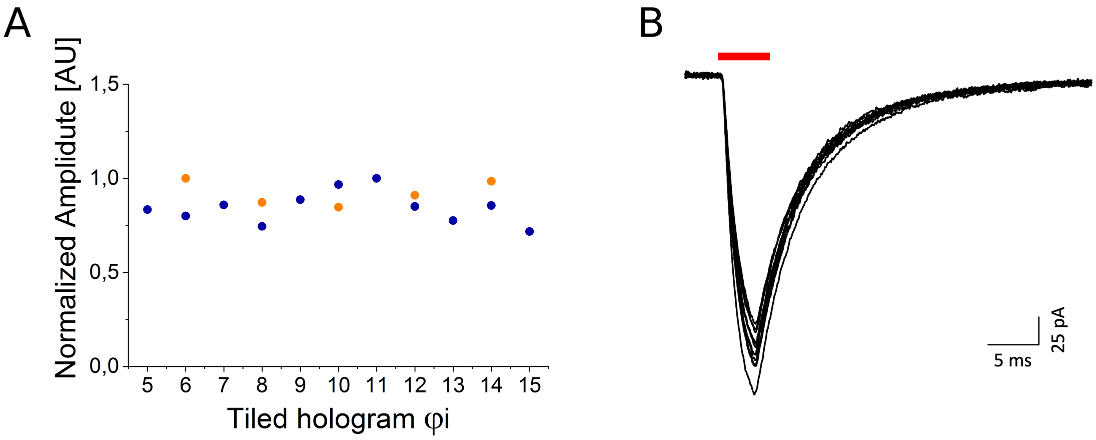


**Supplementary Figure 11:**

**Photocurrents induced by different tiled holograms.**

**(A)** Photocurrent amplitude evoked in the same ST-ChroME-expressing cell by illuminating with a spot encoded by different tiled holograms $\varphi_i$ (n=2 cells). **(B)** Representative photocurrents from a ST-ChroME-expressing patched neuron when illuminated for 5ms with a spot encoded by the tiled hologram $\varphi_{10}$. The SLM was here subdivided in 20 tiled holograms.

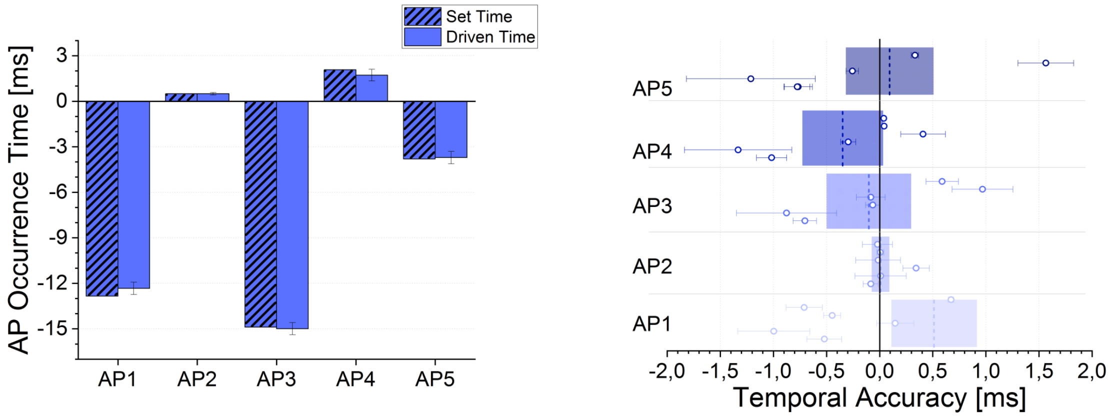


**Supplementary Figure 12:**

**Mimicking of spiking activity.**

AP occurrence time (left) and Temporal precision (right) between imposed and experimental delays of AP peaks for different pairs of double-patched ST-ChroME-expressing cells driven to mimic two independent random patterns of firing as depicted in Fig. 3D. Global mean temporal accuracy is 11 ± 112 µs (n = 12 pair of cells). Mean photostimulation power is 39.8 ± 21.9 mW. Illumination dwell-time ranges between and 2-5ms. 1030nm illumination has been used.

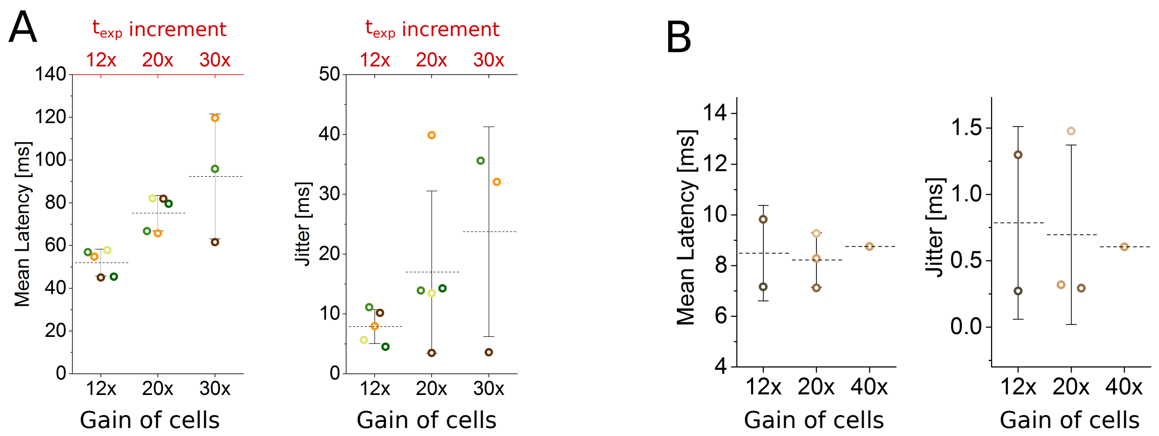


**Supplementary Figure 13:**

**Jitter and latency of light-evoked AP under Multi-S/P cyclic illumination.**

**(A)** Mean latency and jitter of the light-evoked APs obtained in Multi-S/P for different increment of power and total experimental time $t_{exp}$ compared to steady illumination. Different colors indicate different cells. Inset represents threshold power to activate the cells under steady illumination with $t_{std} = 5ms$. Global latency for gain 12x: 50.9 ± 8.6 ms; gain 20x: 76.3 ± 15.4 ms; gain 30x: 95.0 ± 33.6 ms and jitter for gain 12x: 7.9 ± 2.5 ms; gain 20x: 17 ± 12.1 ms; gain 30x: 23.8 ± 14.3 ms. **(B)** Mean latency and jitter of the light-evoked APs obtained in Multi-S/P for different increment of power compared to steady illumination when experimental time $t_{exp}^{cyc}$ = 5ms. Global latency for gain 12x: 8,5 ± 1,5 ms; gain 20x: 8,2 ± 1,1 ms; gain 40x: 8,8 ± 0,5 ms and jitter for gain 12x: 0,8 ± 0,5 ms; gain 20x: 0,7 ± 0,6 ms; gain 40x: 0.6 ms. Different colors indicate different cells. Inset represents threshold power to activate the cells under steady illumination with $t_{std} = 5ms$.

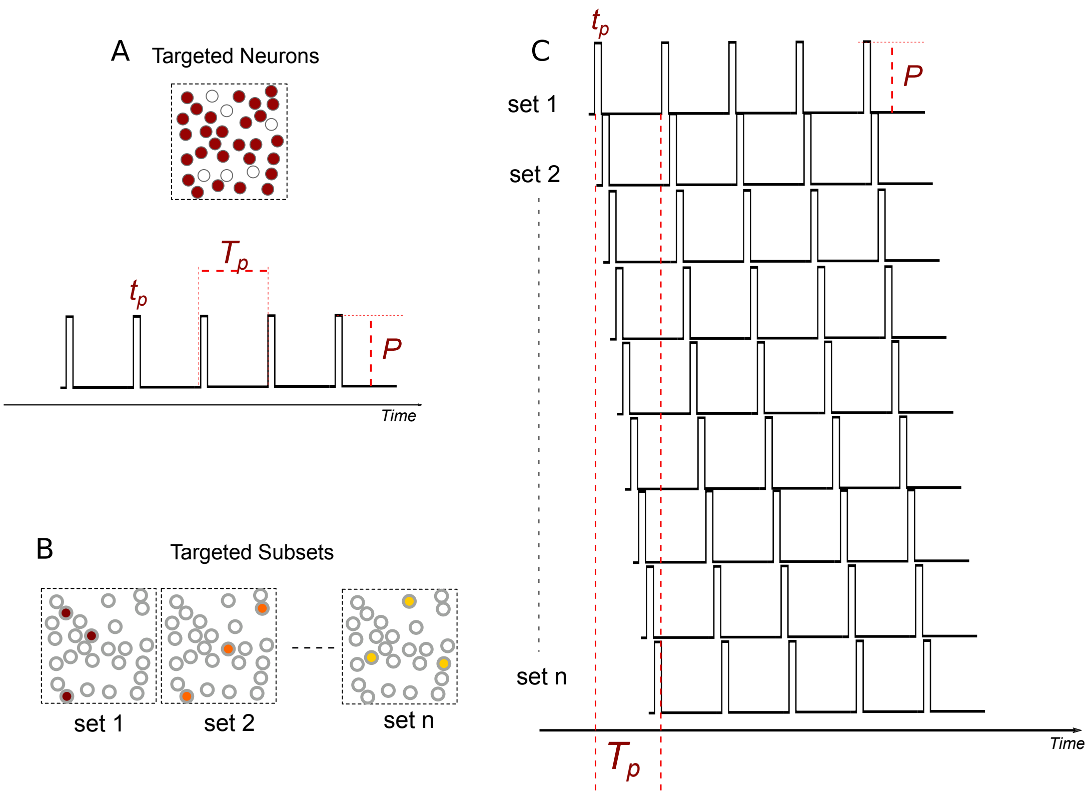


**Supplementary Figure 14:**

**Multi-S/P-FLiT applied to multi-cell activation based on train of ms-range light pulses**.

**(A)** Photostimulation protocol to activate a targeted group of neurons through a train of light pulses of duration $t_p$ and period $T_p$. **(B)** Subsets of the targeted group depicted in (A) encoded in different tiled holograms of an LC-SLM in FLiT configuration. **(C)** Multi-S/P-FLiT illumination protocol resulting by synchronizing the switch between different tiled holograms encoding for different subsets of neurons with the illumination duty-cycle of the pulse train in (A). Different rows correspond to the illumination cyclic occurring on different cell subsets.

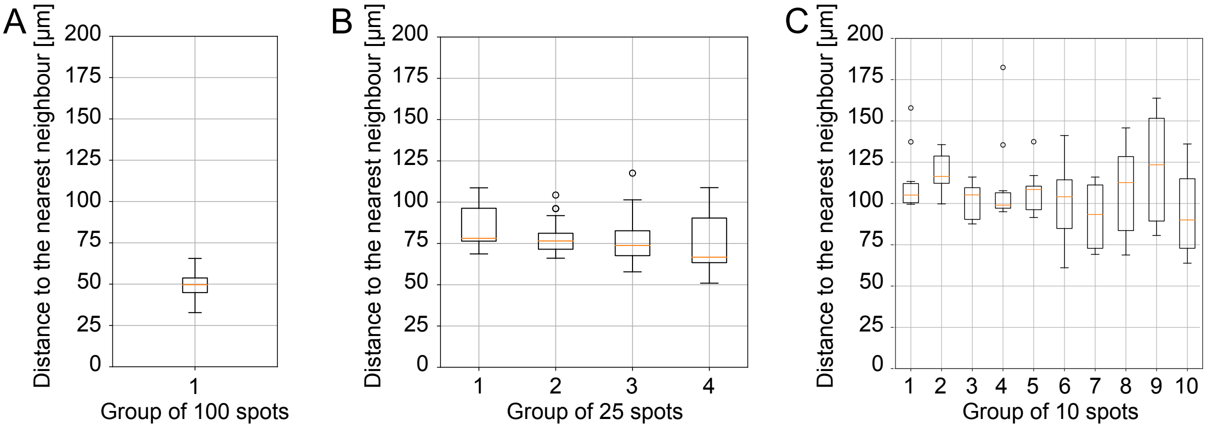


**Supplementary Figure 15:**

**Average distances of spots randomly distributed to maximize the nearest neighbors' distances.**

Average distance to the nearest neighbor for **(A)** 100 spots, **(B)** 10 subsets of 10 spots and **(C)** 4 subsets of 25 spots distributed in a 200x200x500 µm$^3$ volume.

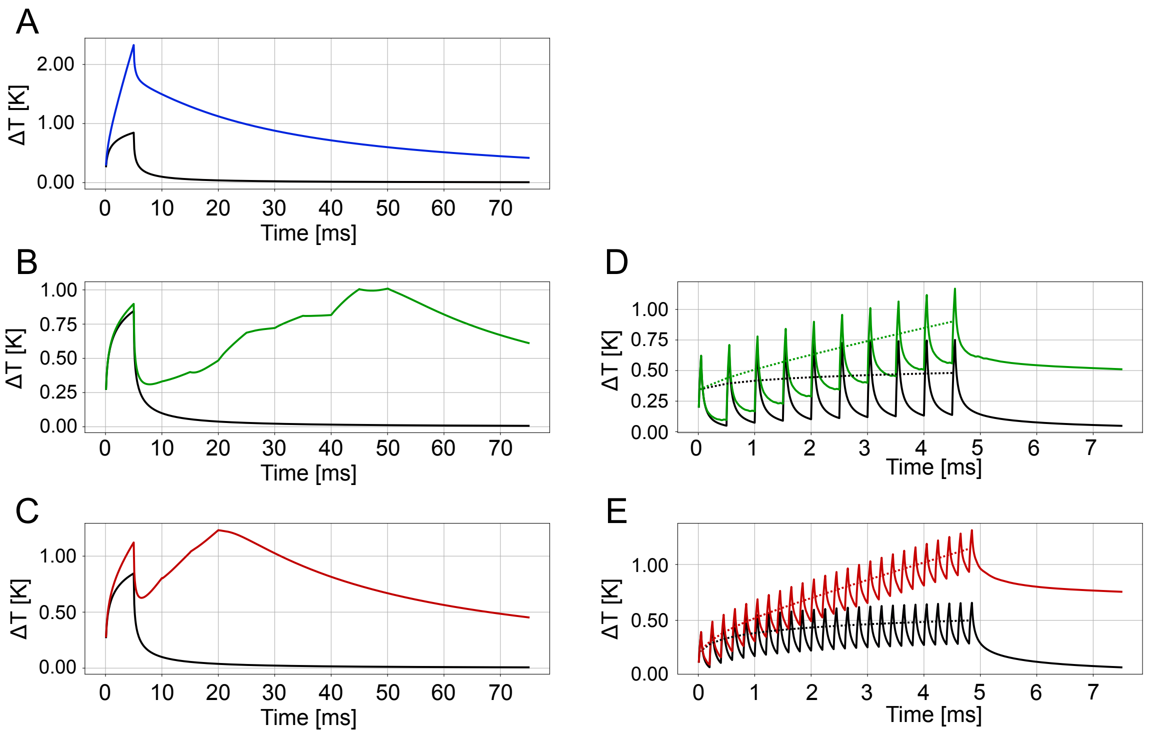


**Supplementary Figure 16:**

**Contribution of single and neighbors' spots to the rise of temperature induced by 100 spots distributed in a 200x200x500 µm$^3$.**

**(A)** Temperature rise averaged on a spot located at coordinate (0,0,0) when the spot in (0,0,0) is illuminated alone with $t_{std} = 5\,ms$ and $P_{std} = 20\,ms$ (black line) or together with 99 neighboring spots randomly distributed in a 200x200x500 µm$^3$ as described in Fig.5, with $t_{std} = 5ms$ and power per spot $P_{std} = 20\,ms$ (i.e., global power P = 2W) (blue line). **(B-C)** Same as A but by sequentially illuminating the 100 spots with n=10 subsets of 10 spots (green line) (B) or n=4 subsets of 25 spots (red line) (C) each illuminated for a time $t_{std} = 5\,ms$. **(D-E)** Same as A but by cyclically illuminating the 100 spots with n=10 subsets of spots for a $t_{exp}$ = 5ms ($t_{cyc}$ = 50 µs; $N_{cyc}$ = 10) and global power per subset $P = 10\sqrt{10} \cdot P_{std}$ (green line) (D) and n=4 subsets of spots for a $t_{exp}$ = 5ms ($t_{cyc}$ = 50 µs; $N_{cyc}$ = 25) and global power per subset $P = 25\sqrt{4} \cdot P_{std}$ (red line) (E). For sequential and cyclic illumination, the spot located in (0,0,0) was part of the first subset illuminated.

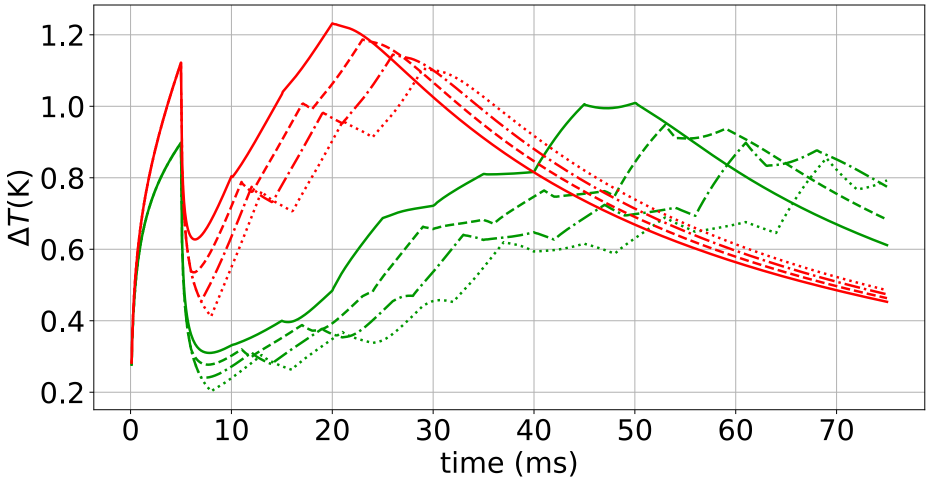


**Supplementary Figure 17:**

**Temperature rise induced by delayed sequential illumination of different subsets of spots.**

Temperature rise averaged on a spot located in (0,0,0) induced by sequentially illuminating 100 spots with n=4 subsets of 25 spots (red lines) or n=10 subsets of 10 spots (green lines) each illuminated for a time $t_{std} = 5\,ms$ when a fixed delay $t_m$ is introduced between the illumination of the different subsets (solid lines $t_m = 0$; dashed lines $t_m = 1ms$; dashed and dots lines $t_m = 2$; dots lines $t_m = 3\,ms$).

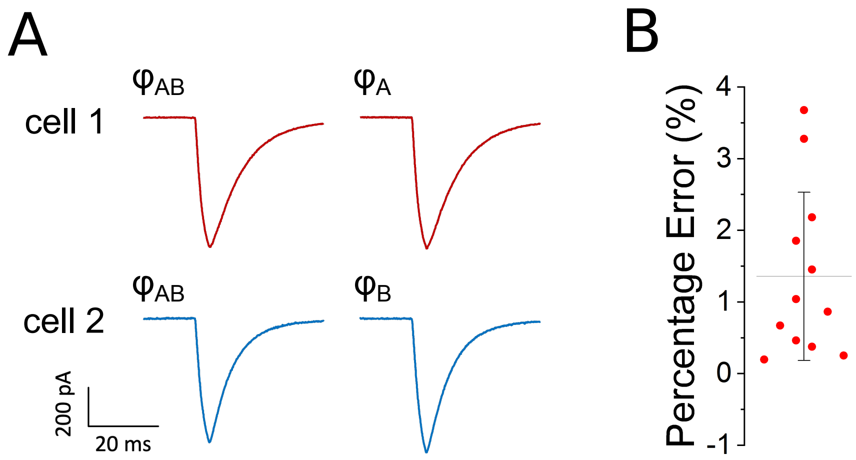


**Supplementary Figure 18:**

**Photo-evoked currents during S/P-FliT experiment.**

**(A)** Representative traces of induced photocurrents in double patched neurons (cell A, red; cell B, blue), when illuminated for 5 ms with a spot encoded by tiled hologram $\varphi_{AB}$ (targeting both cells), $\varphi_{A}$ (targeting cell A) and $\varphi_{B}$ (targeting cell B) and optimizing the illumination power. **(B)** Percentage error of amplitude of induced photocurrents using $\varphi_{AB}$ versus $\varphi_{A}$ or $\varphi_{B}$. Different circle represents different cells. Mean 1.36 ± 1.17 % (n = 12 cells).

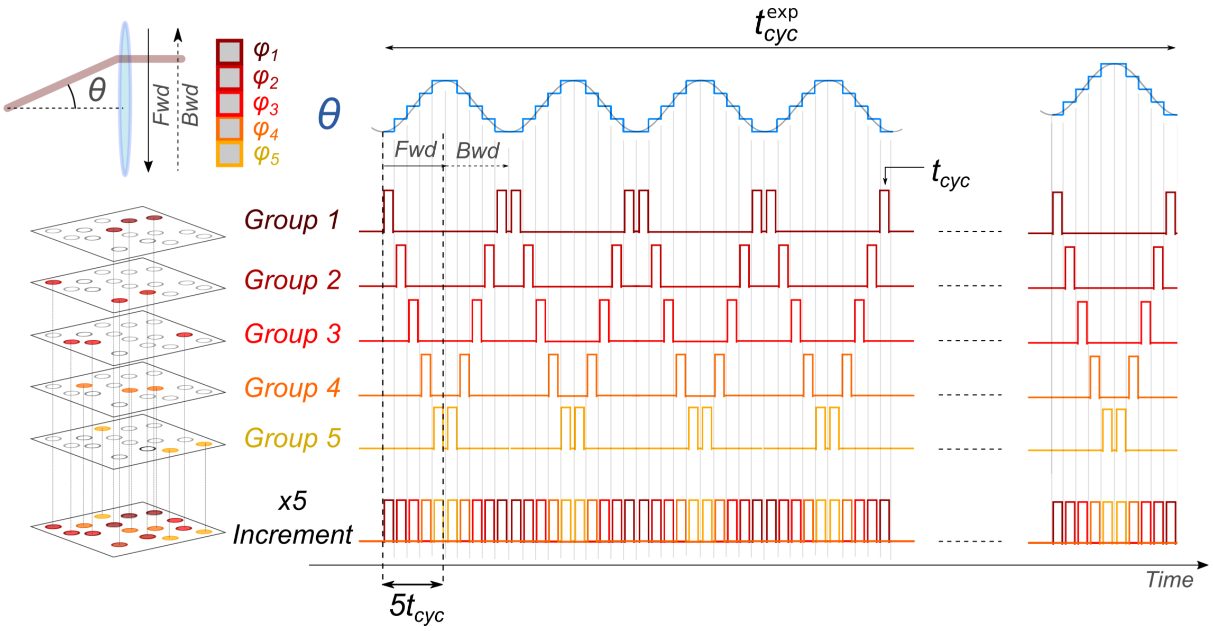


**Supplementary Figure 19:**

**Multi-S/P FLiT illumination protocol.**

The LC-SLM is tiled in *n* different tiled holograms $\varphi_i$, each encoding for different groups of cells (in the present scheme from 1 to 5). The laser is continuously steered back and forward across the LC-SLM by varying the deflection angle $\theta$ of the galvanometric mirror following a staircase input voltage. Each hologram is illuminated for a dwell time $t_{cyc}$ over one cycle whose duration correspond to $n \cdot t_{cyc}$ (in the present scheme corresponding to $5t_{cyc}$) for a total duration $t_{exp}^{cyc}$. The scheme displayed is meant to represent *n* groups of spots; their number is here limited to 5 for presentation purposes only.

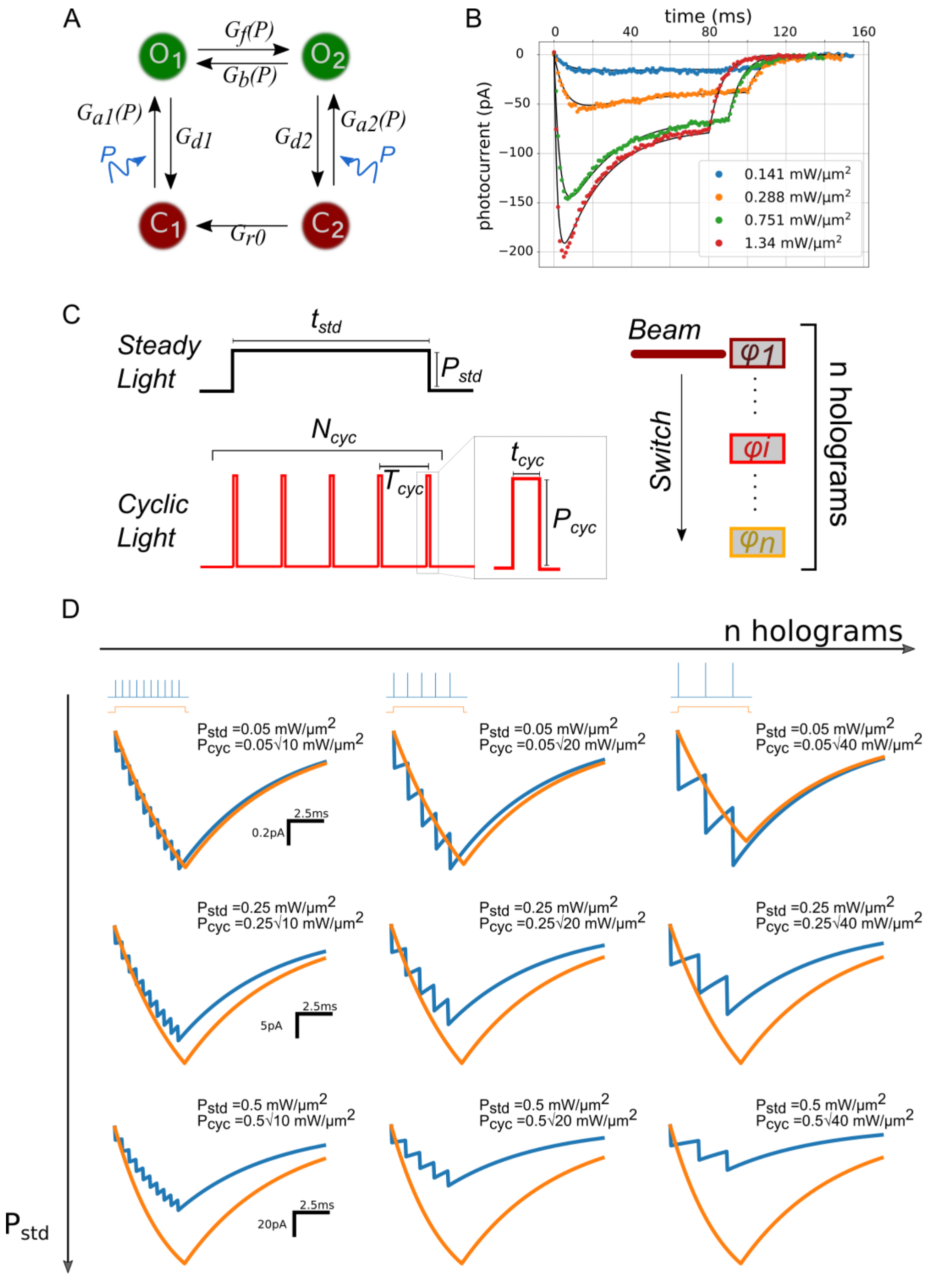

**Supplementary Figure 20: Simulated photocurrents under steady and cyclic illumination. (A)** Scheme of four-state model with $G_{a1}(P)$, $G_{a2}(P)$, $G_{d1}$, $G_{d2}$, $G_f(P)$, $G_b(P)$ and $G_r$ indicating the rate constants for transitions $C_1 \rightarrow O_1$, $C_2 \rightarrow O_2$, $O_1 \rightarrow C_1$, $O_2 \rightarrow C_2$, $O_1 \rightarrow O_2$, $O_2 \rightarrow O_1$ and $C_2 \rightarrow C_1$, respectively. **(B)** Fitting of experimental photocurrent traces obtained by photostimulating a ST-ChroME-expressing neuron for different light intensities. **(C)** Scheme of photostimulation under steady illumination of power $P_{std}$ and duration $t_{std}$ and under cyclic illumination of power $P_{cyc}$ and pulse duration $t_{cyc}$ over $N_{cyc}$ cycles. **(D)** Simulated photocurrents of a light-targeted neuron in the approximation of a four-state model under 2PE steady illumination (orange) or cyclic illumination with $P_{cyc} = P_{std}\sqrt{n}$ (blue) for different values of illumination power and number of holograms $n = T_{cyc}/t_{cyc}$.

**SUPPLEMENTARY MOVIES:**

**Supplementary Movie 1: Alternation of different groups of spots in FLiT.**

Different groups of spots are alternated by sequentially tilting the galvanometric mirror on different tiled holograms $\varphi_i$ of the multiplexing LC-SLM, each encoding for different 2D patterns. The LC-SLM was subdivided into 10 different tiled holograms. The tiled holograms were updated after scanning the entire LC-SLM by automatically refreshing the LC-SLM with new phase masks. FoV is 150x150 $\mu m^2$. Time per frame 20 ms.

**Supplementary Movie 2: Time lapse of the 3D distribution of the temperature rise during sequential illumination with *n* subsets of spots.**

**From left to right:** Temperature rise induced by steadily illuminating one set of 100 spots in parallel as shown in Fig. 5C (blue line) with $t_{exp} = t_{std}$=5ms and global power $P = 100 \cdot P_{std}$ (with $P_{std}$ =20mW); Temperature rise induced by steadily illuminating the 100 spots in sequence with n=4 subsets of spots as shown in Fig.5C (red line) with $t_{exp}$=20ms and global power per subset $P = 25 \cdot P_{std}$; Temperature rise induced by steadily illuminating the 100 spots in sequence with n=10 subsets of spots as shown in Fig. 5C (green line) with $t_{exp}$=50ms and global power per subset $P = 10 \cdot P_{std}$; Temperature rise induced by illuminating one spot individually under steady illumination for $t_{exp} = t_{std}$=5ms as shown in Fig.5C (black line).

**Supplementary Movie 3: Time lapse of the 3D distribution of the temperature rise during cyclic illumination with *n* subsets of spots.**

**From left to right:** Temperature rise induced by cyclically illuminating 100 spots as shown in Fig.5D (red line) with n=4 subsets of spots for a $t_{exp}$=5ms ($t_{cyc}$=50µs; $N_{cyc}$=25) and global power per subset $P = 25\sqrt{4} \cdot P_{std}$; Temperature rise induced by cyclically illuminating 100 spots as shown in Fig.5D (green line) with n=10 subsets of spots for a $t_{exp}$=5ms ($t_{cyc}$=50µs; $N_{cyc}$=10) and global power per subset $P = 10\sqrt{10} \cdot P_{std}$; Temperature rise induced by illuminating one spot individually.
